# Contrasting effects of *Ksr2*, an obesity gene, on trabecular bone volume and bone marrow adiposity

**DOI:** 10.1101/2022.08.21.504689

**Authors:** Gustavo A. Gomez, Charles H. Rundle, Weirong Xing, Chandrasekhar Kesavan, Sheila Pourteymoor, Robert E. Lewis, David R. Powell, Subburaman Mohan

## Abstract

Pathological obesity and its complications are associated with an increased propensity for bone fractures. Humans with certain genetic polymorphisms at the kinase suppressor of ras2 (*Ksr2*) locus develop severe early-onset obesity and type 2 diabetes (T2D). Both conditions are phenocopied in mice with *Ksr2* deleted, but whether this affects bone health remains unknown. Here we studied the bones of global *Ksr2* null mice and found that *Ksr2* negatively regulates femoral, but not vertebral, bone mass in two genetic backgrounds, while the paralogous gene, *Ksr1*, was dispensable for bone homeostasis. Mechanistically, KSR2 regulates bone formation by influencing adipocyte differentiation at the expense of osteoblasts in the bone marrow. Compared with *Ksr2*’s known role as a regulator of feeding by its function in the hypothalamus, pair feeding and osteoblast-specific conditional deletion of *Ksr2* reveals that *Ksr2* can regulate bone formation autonomously. Despite the gains in appendicular bone mass observed in absence of *Ksr2*, bone strength, as well as fracture healing response remains compromised in these mice. This study highlights the interrelationship between adiposity and bone health and provides mechanistic insights into how *Ksr2*, an adiposity and diabetic gene, regulates bone metabolism.

## Introduction

Obesity is a major public health problem in the Unites States, afflicting nearly 40% of adults and has become a prevalent and destructive health disorder linked to some of the major metabolic diseases including cardiovascular diseases, type 2 diabetes (T2D), and cancer (Devlin and Rosen, 2015; Pagnotti et al., 2019; Shanbhogue et al., 2016; Walsh and Vilaca, 2017). Although obesity may be considered beneficial to bone health, since increased body weight is associated with higher bone mineral density (BMD), the relationship between excess body fat and bone health is complex, given that obesity has been identified as a risk factor for certain fractures (Greco et al., 2015; C. Ma et al., 2018; Veldhuis-Vlug and Rosen, 2018). The increasing prevalence of obesity and T2D dictates the need for appropriate in vivo animal models to study the mechanisms of action of obesity and T2D on bone metabolism.

The effect of obesity and T2D on bone is an active area of investigation. Studies with several animal models and approaches have contributed to our current understanding of this relationship. Mouse models, in particular, have provided invaluable information through controlled genetic, biochemical, cellular, and molecular approaches to understand the pathological relationship between excess body fat and bone fragility. Most diet-induced obesity studies have reported reduced BMD and trabecular bone mass (Bonnet et al., 2014; Doucette et al., 2015; Inzana et al., 2013; Scheller et al., 2016). By contrast, monogenetic models of obesity have provided a broader range of bone phenotypes including no change, loss, or gain in bone mass or BMD (Ahn et al., 2006; Baldock et al., 2007; Braun et al., 2012; Steppan et al., 2000; X. Wang et al., 2007). There are several explanations for the diversity in skeletal phenotypes in these models including differences in expression of targeted genes in other tissues besides bone as well as varied effects of endocrine factors produced in other affected tissues such as the brain, fat, and skeletal muscle. Nevertheless, monogenetic studies have informed the molecular underpinnings of feeding regulation at the hypothalamus, which has fortuitously led to the development of pharmaceuticals to treat a particular population of individuals genetically predisposed to diabetes (Yeo et al., 2021). Although the RANKL monoclonal antibody, denosumab, has been used to treat bone disorders in osteoporotic T2D patients with reduced BMD (Abe et al., 2019), whether these agents can also benefit the population with gains in BMD, which are paradoxically also fragile (Burghardt et al., 2010; L. Ma et al., 2012), remains to be investigated. Also, it is worthwhile to further identify/study animal models with genetic mutations that phenocopy the human condition to study these interventions. The advent of the genomic era has expanded the list of individual genes associated with obesity and T2D (Loos and Yeo, 2022), yet their effects on bone remain vastly understudied.

Recently, the scaffold proteins kinase suppressor of ras (KSR1 and KSR2) were identified as mediators of energy consumption, utilization, and adipogenic regulation (Brommage et al., 2008; Costanzo-Garvey et al., 2009; Kortum et al., 2005; Pearce et al., 2013; Revelli et al., 2011). Although these two genes function as paralogs, we previously found that only *Ksr2* knockout (KO) mice become obese and diabetic (Brommage et al., 2008; Costanzo-Garvey et al., 2009; Kortum et al., 2005; Pearce et al., 2013; Revelli et al., 2011), suggesting these paralogs have non-redundant roles, although *Ksr1* does have a role in adipogenesis (Kortum et al., 2005). Several mutations at the *Ksr2* loci in humans have been associated with severe early-onset obesity (Pearce et al., 2013), and studies in *Ksr2* KO mice have revealed a centrally regulated mechanism by *Ksr2* expression and function in the hypothalamus that results in hyperphagia, changes in metabolic rate, and consequently, obesity and T2D (Costanzo-Garvey et al., 2009; Guo et al., 2017; Henry et al., 2014; Pearce et al., 2013; Revelli et al., 2011). Although there are hundreds of mouse genes reported to lead to obesity when disrupted, *Ksr2* gene deletion is associated with a profound obese phenotype and lethality at a young age (Brommage et al., 2008; Revelli et al., 2011). In this study, we set out to investigate whether the deletion of *Ksr2*, an obesity and T2D gene, bears any effect on bone health and if so, to evaluate the mechanisms by which KSR2 affects bone metabolism. Our studies show that loss of KSR2 function increases long bone trabecular bone mass while reducing marrow adiposity and that KSR2 acts as a molecular switch that controls the differentiation of bone resident mesenchymal stem cells into osteoblast or adipocyte differentiation via an mTOR-dependent mechanism.

## Results

### Ksr2 negatively regulates femoral bone mass

To evaluate whether deletion of *Ksr2* affects skeletal morphology, femurs of *Ksr2* KO (exon 13 deleted) and wild-type (WT) control mice in the C57BL/6J-Tyr*^c-Brd^* X 129^SvEvBrd^ hybrid background (Figure 1A), were subjected to microCT scanning. Distal femoral metaphyseal bones of *Ksr2* KO (*Ksr2^-/-^*) female mice exhibited increased trabecular bone mass at both 11 and 15 weeks of age (Figure 1B). Quantification of trabecular parameters at the distal femur secondary spongiosa shows that by 11 weeks, bone volume fraction (BV/TV), and trabecular thickness (Tb. Th) were significantly increased in female KO mice (Figure 1C, G), while structure model index (SMI), a measure of rod to plate-like trabecular morphology (Hildebrand and Rüegsegger, 1997), was significantly reduced (Figure 1E), overall implying that structural morphological changes elicited by the absence of *Ksr2* promotes gains in bone mass. Although mean trabecular connectivity density (CONN.D), and trabecular number (Tb.N) were also increased in KO mice at 11 weeks (Figure 1D and F), these differences were not significant until 15 weeks, as was the reduction in trabecular spacing (Figure 1H).

**Figure 1.**
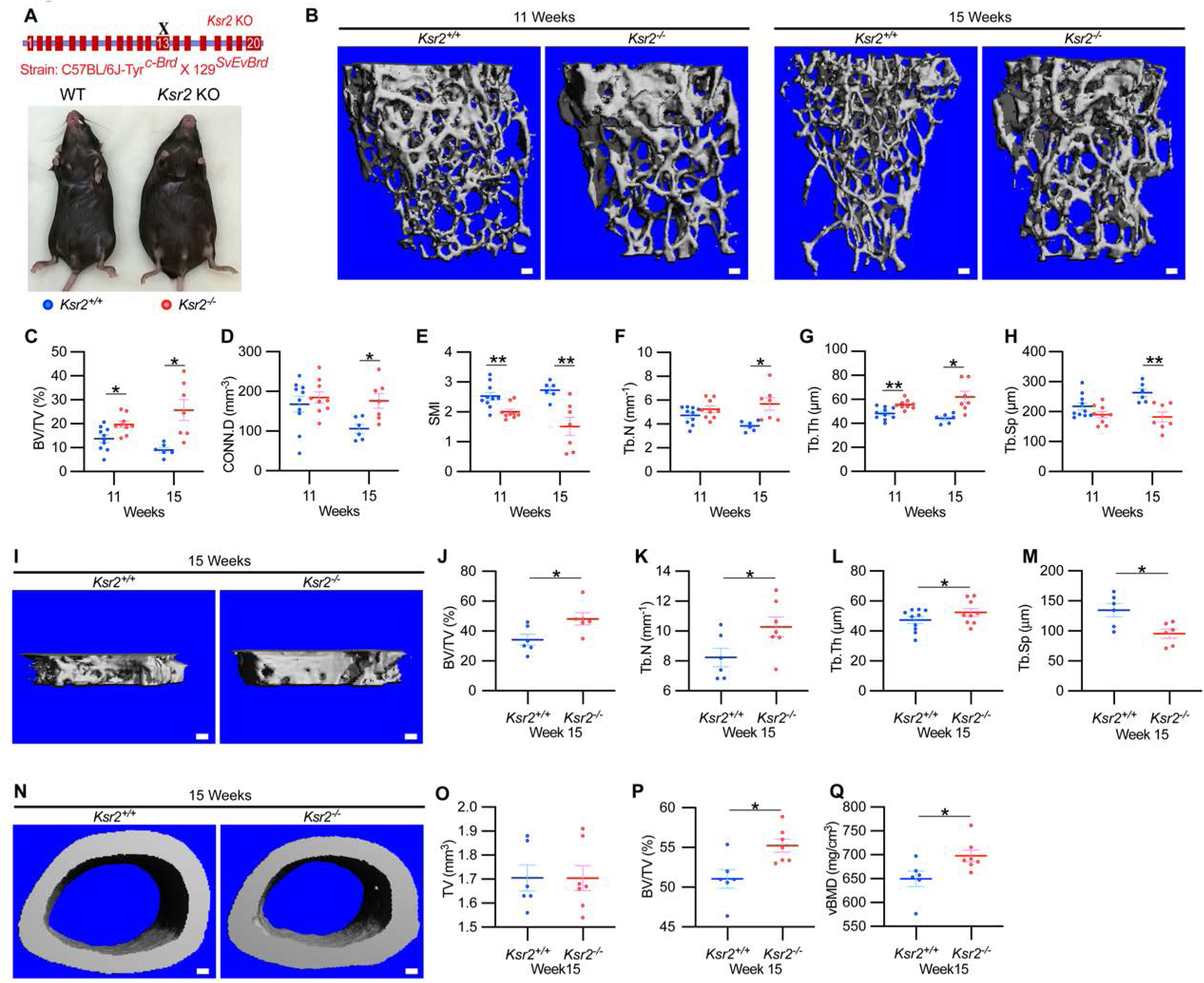
*Ksr2* regulates bone mass in females. (A) Schematic of *Ksr2* knocked out in the C57BL/6J-Tyr*^c-Brd^* X 129^SvEvBrd^ hybrid strain with exon 13 deleted (X), and accompanying ventral view of genotyped mice at 4 months of age exhibiting differences in weight gain. (B) Representative 3D microCT reconstruction images of the secondary spongiosa at the distal femoral metaphysis in wild-type (*Ksr2^+/+^*, WT) or knockout (*Ksr2^-/-^*, KO) females at 11 and 15 weeks, revealing a prominent increase in trabecular bone in KOs. Scale bar: 100μm. (C-H) MicroCT measurements from the trabecular bone as represented in part B (n = 6-10/group), Abbreviations: BV/TV, Bone Volume/Tissue Volume; CONN.D, Connectivity Density; SMI, Structural Model Index; Tb.N, Trabecular number; Tb.Th, Trabecular thickness; Tb.Sp, Trabecular spacing. (I) Representative 3D reconstruction of microCT images of primary spongiosa in WT or KO mice at 15 weeks of age revealing increased bone density in KO mice. Scale bar: 100μm. (J-M) Quantification of microCT parameters measured in part I (n = 6-10/group). (N) Representative 3D reconstruction of microCT images of cortical bone at the femoral mid-diaphysis (Scale bar: 100μM), where the TV total volume (O) is not affected, while BV/TV and vBMD volumetric Bone Mineral Density (P, Q) are increased in KO mice. Statistics analyzed by unpaired 2-tailed student t-test, and graphed lines represent the mean ± SEM, * p<0.05, ** p<0.005

New woven bone is actively formed and mineralized at the primary spongiosa while the woven bone is remodeled into mechanically stronger lamellar bone at the secondary spongiosa. To determine if new bone formation at the primary spongiosa is altered in the *Ksr2* KO mice, we measured trabecular bone parameters at the primary spongiosa, limited to within 300μm of the distal most femoral metaphyseal bone from the growth plate (Figure 1I-M) and found significant increases in trabecular bone volume. *Ksr2* deleted males also exhibited significantly greater BV/TV, Conn. Den, Tb. N and Tb. Th but reduced Tb. Sp and SMI compared to littermate control mice (Figure 2A-G). Thus, loss of the *Ksr2* gene promotes trabecular bone density in both genders of mice.

**Figure 2.**
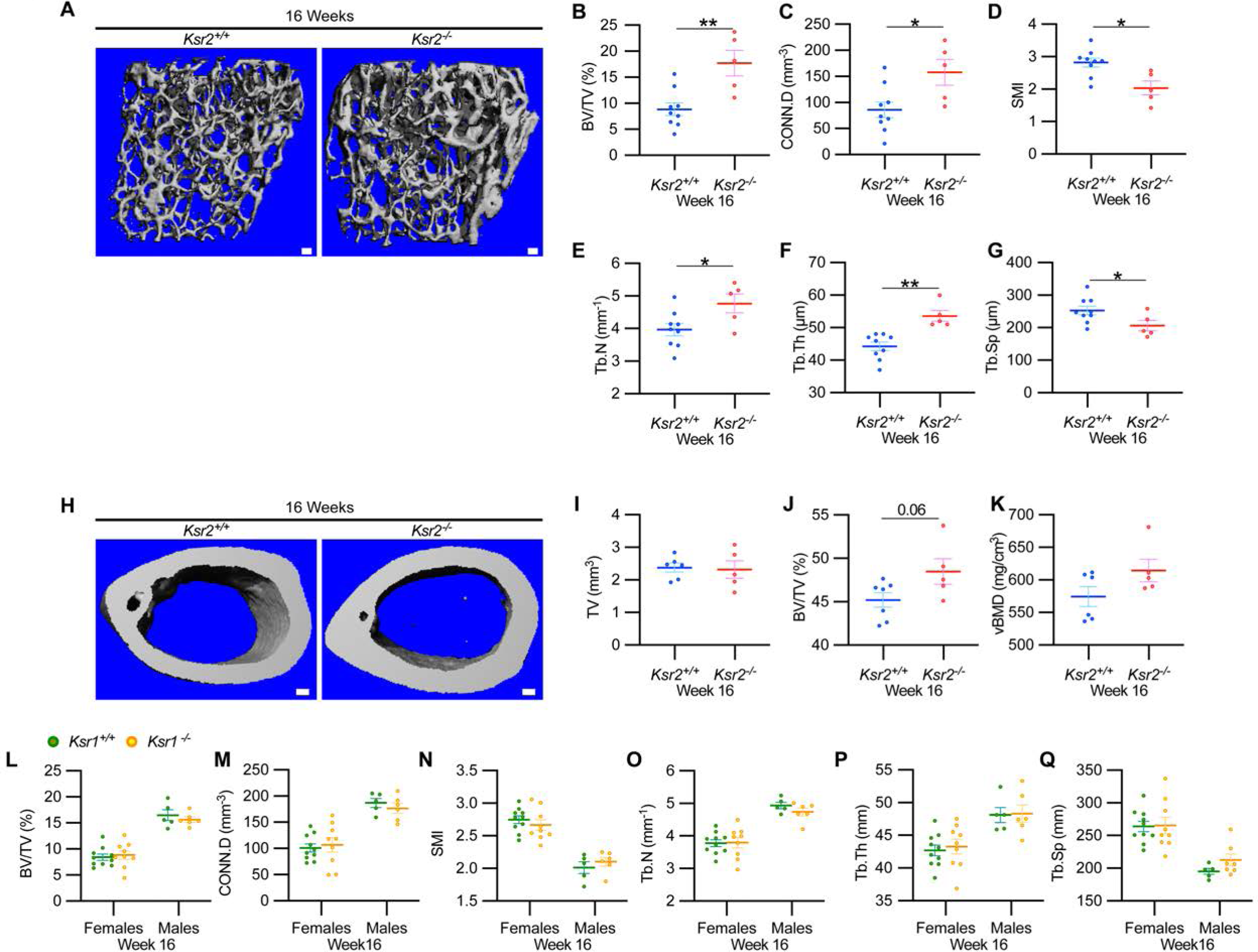
*Ksr2* negatively regulates femoral bone in males, while *Ksr1* deletion does not affect trabecular bone in either gender. (A) Representative 3D microCT reconstruction images of the distal femoral metaphysis in WT or KO male mice at 16 weeks of age revealing increased trabecular bone in KOs. Scale bar: 100μm. (B-G) MicroCT measurements from the trabecular bone as represented in part A (n = 5-9 mice per group). Abbreviations: BV/TV, Bone Volume/Tissue Volume; CONN.D, Connectivity Density; SMI, Structural Model Index; Tb.N, Trabecular number; Tb.Th, Trabecular thickness; Tb.Sp, Trabecular spacing. (H) Representative 3D microCT reconstruction images of cortical bones at the femoral mid-diaphysis revealing that *Ksr2* deletion does not affect TV total volume (I) of cortical bone in males, while BV/TV Bone volume/tissue volume (J)and vBMD volumetric bone mineral density (K) are increased in Ksr2 KO mice (n = 5-6/group). Scale bar: 100μm. (L-Q) Quantification of microCT data from the distal femoral metaphysis of WT and *Ksr1* knockout mice at 16 weeks of age, showing minimal changes in trabecular bone parameters between genotypes in either gender. Statistics analyzed by 2-tailed student t-test, and graphed lines represent the mean ± SEM, * p<0.05, ** p<0.005

By further characterization of femoral bones, we found increased mid-shaft femoral cortical bones in *Ksr2* nulls. While total tissue volume, indicative of bone size, remained unchanged in the KO mice in both genders (Figure 1N-P, Figure 2H-K), a significant increase in bone volume fraction was observed, although gains in bone mineral density were more prominent in females. Nonetheless, this evaluation reveals that obese *Ksr2* null mice present increased gains in trabecular and cortical mass of femoral bones.

### Ksr1 is dispensable for the development of femoral trabecular bone mass

*Ksr1*, the only paralog of *Ksr2*, is highly expressed in skeletal muscle (Costanzo-Garvey et al., 2009), which interacts with, and affects bone physiology (Lara-Castillo and Johnson, 2020). *Ksr1* was also expressed in osteoblasts (data not shown). To determine whether *Ksr1* might also contribute to limb bone mass accretion, we evaluated metaphyseal femoral bones of 16-week-old *Ksr1* KO mice and their wild-type littermate controls by microCT. In contrast with the striking differences found in trabecular bone parameters in *Ksr2* KO mice at this age, trabecular bone measurements were nearly identical between *Ksr1* KO and WT mice, irrespective of gender (Figure 2, L and M). These results suggest that the KSR1 protein is highly unlikely to synergize with KSR2 in regulating femoral bone growth.

### Validation of bone phenotype by Ksr2 deletion in a different genetic background

Since genetic background can influence biological effects in mice (Ackert-Bicknell et al., 2009; Bonnet et al., 2014), we evaluated whether *Ksr2* deleted in the DBA/1LacJ strain, which also becomes obese (Costanzo-Garvey et al., 2009), might also exhibit alterations in skeletal phenotype. Deletion of exon 4 in this genetic background did not significantly affect the anus to nose body length, although a substantial gain in body weight, and percent body fat was observed by 8 weeks of age (Figure 3A-D). Concordant with *Ksr2* deletion in the C57BL/6J-Tyr*^c-Brd^* X 129^SvEvBrd^ hybrid background, dual-energy X-ray absorptiometry (DXA) measurements of pooled genders revealed substantial increases in total body bone mineral density (Figure 3E) and in particular, in femoral bone mineral density (Figure 3G), while femur length remained unchanged relative to WT siblings (Figure 3F). Additionally, no changes were observed in the vertebral trabecular bone (Figure 3—figure supplement 1).

**Figure 3.**
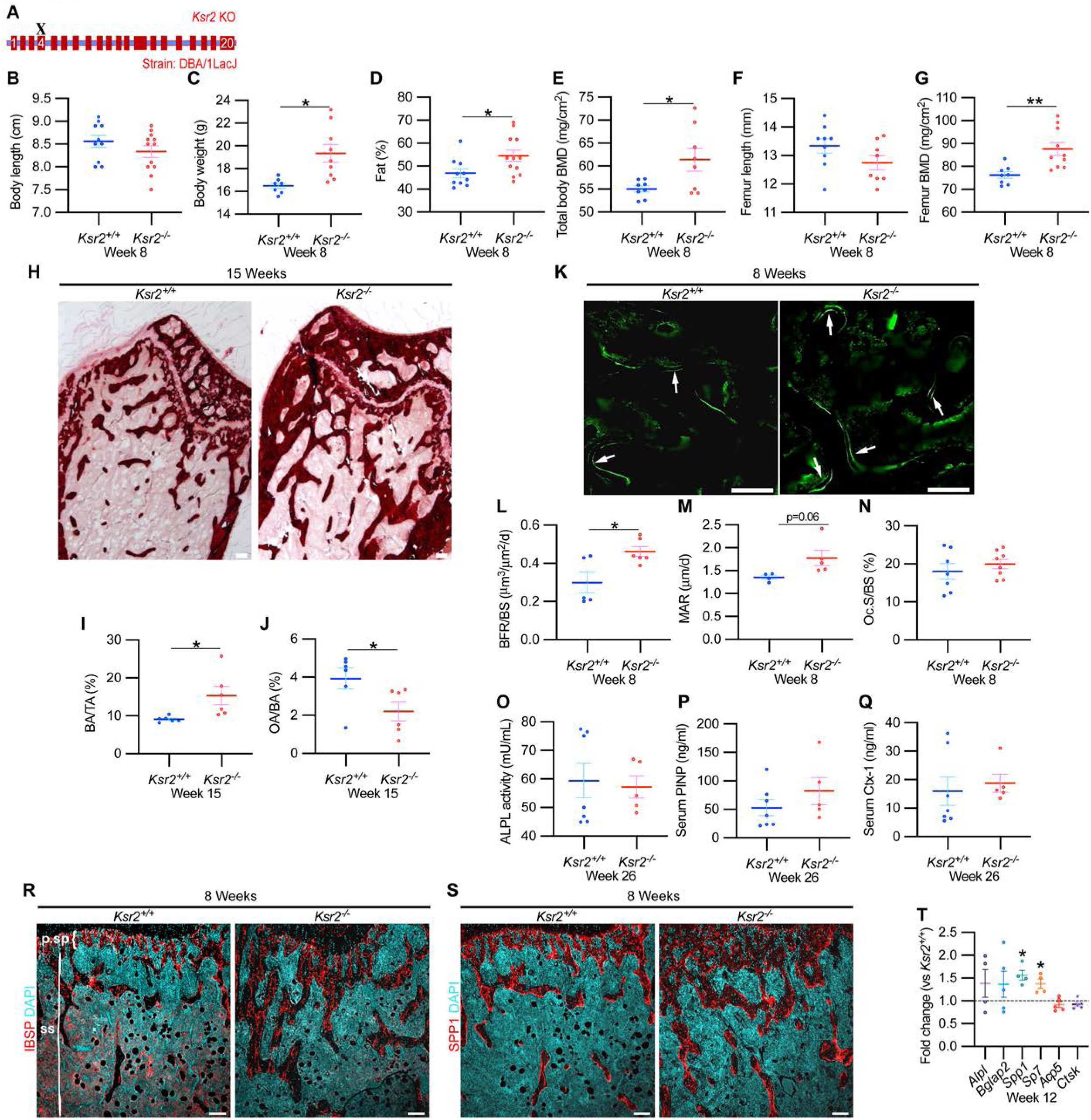
*Ksr2* deletion in a different genetic background, histomorphometry, and histology validate that *Ksr2* negatively regulates bone formation. (A) Schematic of *Ksr2* knocked out in the DBA/1LacJ strain with exon 4 deleted (X). (B) No differences noted in body length at 8 weeks of age, while gains in body weight (C) and body fat percentage (D) are noted in KOs. BMD is increased in total body (E) and femurs (G) of KO mice, while femur length is not changed (F) (n = 7-12 mice/group; genders combined), (D-G reflect dual-energy x-ray absorptiometry measurements). (H) Representative Alizarin red images at the distal femoral epiphysis show increased area of mineral staining in KO mice at 11 weeks of age. Scale bar: 100μm. (I,J) Quantification of Alizarin stain reveals an increase in BA/TA, bone area/total area, and a decrease in OA/BA, osteoid area/bone area. (K) Representative histomorphometric images of fluorescent calcein label reveal increased staining in KO mice. Scale bar: 100μm. (L-N) Quantification of histomorphometric parameters measured, showing increased bone formation rate/bone surface, BFR/BS, and Mineral apposition rate, MAR, yet no changes in number of osteoclasts per bone surface, Oc.S/BS. (n = 4-7 mice/group). (O-Q) No changes in ALPL, PINP, and Ctx-1 activity in serum at 26 weeks (n=5-7 mice/group). (R,S) Immunofluorescence staining at distal femoral metaphysis for (IBSP, synonym BSP2) or (SPP1, synonym OPN) (both red), counterstained with DAPI (cyan) reveals broader expression of both bone markers in KO mice; growth plate-osteoblast boundary positioned at the top. Abbreviations: p.sp, primary spongiosa; ss, secondary spongiosa. Scale bar: 100μm. (T) RT-qPCR reveals increased expression of osteoblast markers (*Alpl*, *Bglap2*, *Spp1* and *Sp7*), while osteoclast markers (*Acp5*, *Ctsk*) remain unchanged in femurs of KO mice. Statistics analyzed by 2-tailed student t-test, and graphed lines represent the mean ± SEM, * p<0.05, ** p<0.005

Histological evaluation of longitudinal distal metaphyseal femur bone sections by alizarin red also revealed increased amounts of calcified bone in *Ksr2* KO mice, as the ratio of bone area over total area was increased, while osteoid area over bone area was decreased (Figure 3H-J, Figure 3—figure supplement 2). Moreover, qualitative comparisons of bone markers, Integrin bone sialoprotein (IBSP) and Secreted phosphoprotein 1 (SPP1) by immunofluorescence suggest an increased areal expansion of both markers throughout the metaphysis in KO mice (Figure 3, R and S). Overall, this data provides further supporting evidence that *Ksr2* negatively regulates appendicular bone formation, with confirmation in a different genetic background.

### Gains in bone mass in Ksr2 nulls are a product of increased osteoblast activity

Next, we began to address how the deletion of *Ksr2* results in increased bone mass. To determine if increased bone formation is the cause of increased bone mass in *Ksr2* KO mice, we performed histomorphometric analysis by pulsed calcein injections in 8-week-old mice. This resulted in increased calcein labeling in KO mice, with quantitative gains detected in bone formation rate and mineral apposition rate (Figure 3K-M). By contrast, the percentage of bone-resorbing acid phosphatase 5, tartrate resistant (ACP5+) osteoclasts scored per bone surface did not change (Figure 3N). However, serum comparisons of markers indicative of osteoblast bone deposition (Alkaline phosphatase, ALPL; and procollagen type 1 N-terminal propeptide, P1NP) or osteoclast activity (Carboxy-terminal cross-linked telopeptide of type 1 collagen, CTX1) revealed no differences in ALP activity or CTX1 levels between the two genotypes at 6 months of age (Figure 3O-Q).

Bulk comparisons between osteoblast and osteoclast markers were then compared by real time-quantitative PCR (RT-qPCR) from the metaphysis of *Ksr2* KO relative to WT littermates at 12 weeks of age. mRNA expression levels of osteoblast markers, *Alpl* and *Bglap2* showed increased, though insignificant, expression in KO mice, while *Spp1* and *Sp7* were significantly increased. By contrast, markers of differentiated osteoclasts, *Acp5* and *Ctsk*, remained unchanged (Figure 3T). Combined, these results posit that *Ksr2* affects osteoblast function, and does not apparently affect osteoclasts.

### Ksr2 gains bone at the expense of adipocyte differentiation

As *Ksr2* KO mice are obese, exhibiting increased visceral and subcutaneous adiposity (Figure 4A), we determined if genetic disruption of *Ksr2* influences adipocyte gene expression in white and brown adipose tissues in 28-week-old mice. mRNA levels of key transcription factors, *Pparg* and *Cebpa* were unchanged in both fat depots in *Ksr2* KO mice (Figure 4B) at this age. By contrast, the adipokine leptin (*Lep*) was increased, while complement factor D (*Cfd*) was decreased in both white and brown fat of *Ksr2* KO mice (Figure 4C). This suggests that KSR2 exerts opposite effects on leptin and complement factor D/adipsin expression in fat tissues, which is consistent with changes observed in other models of obesity in mice (Cianflone et al., 2003; Kwon et al., 2012).

**Figure 4.**
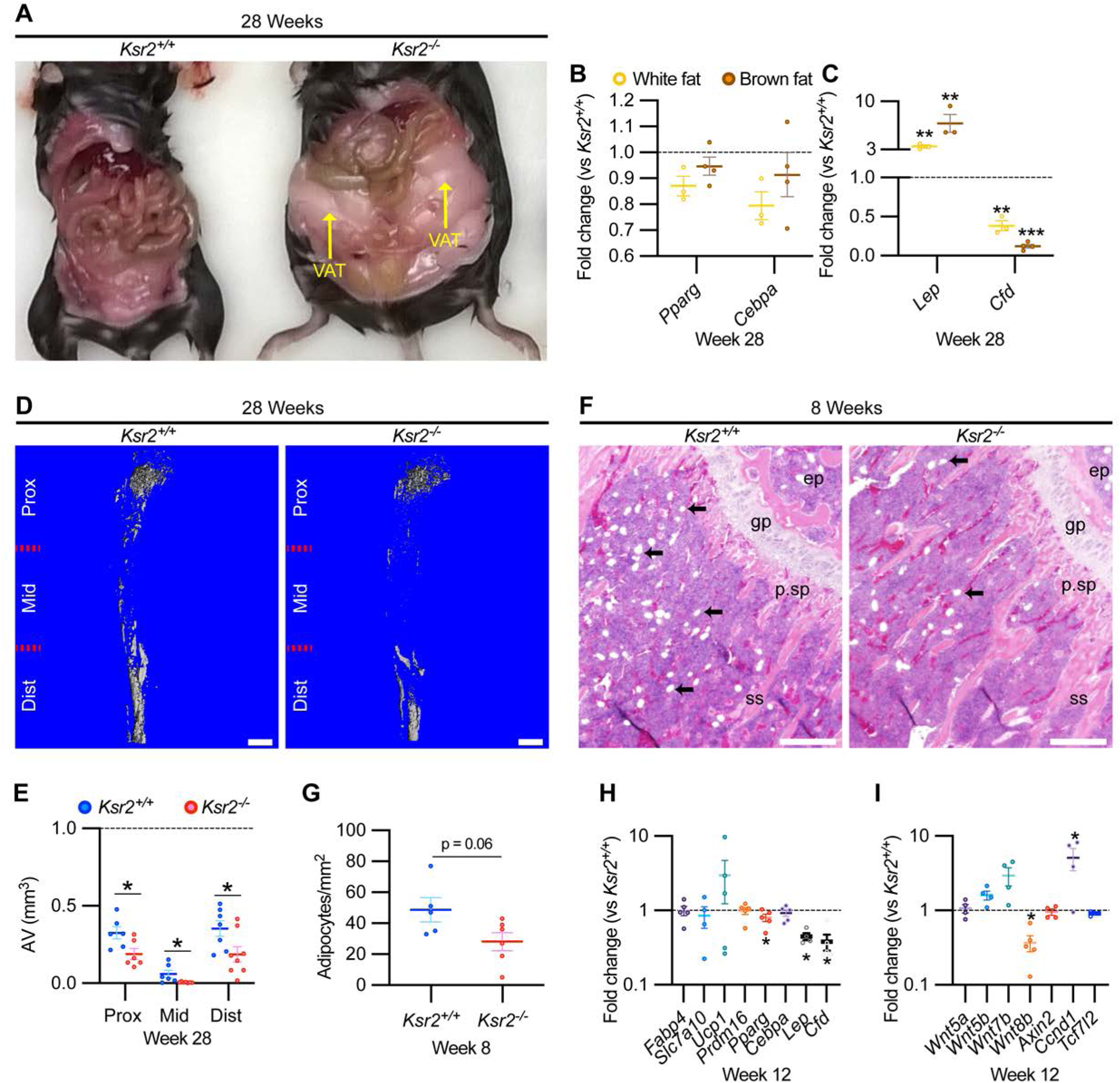
Obesity in *Ksr2* null mice paradoxically presents reduced bone marrow adiposity. (A) Representative image of mice at 28 weeks of age dissected to reveal differences in visceral adipose tissue (VAT) between WT and *Ksr2* KOs. (B-C) RT-qPCR assessing changes in regulators of adipogenesis (B) or adipokine genes (C), in white or brown fat of *Ksr2* KO mice relative to WT at 28 weeks of age (n = 3-4/group). (D) Representative 3D microCT reconstruction images of osmium tetroxide labeled femurs, revealing reductions in bone marrow adipose tissue in *Ksr2* KO mice at 28 weeks of age. Scale bar: 1mm. (E) Quantification of adipocyte volume (AV) occupied by marrow adipose tissue in femurs of mice as depicted in part D, at proximal (prox), middle (mid), and distal (dist) thirds of the femur with position defined in reference to the spinal cord. (n = 6-8/group). (F) Representative hematoxylin and eosin-stained longitudinal distal femur sections of 8-week-old mice in which adipocytes (arrows) were compared at the secondary spongiosa, revealing reductions in KO mice. Scale bar: 100μm. (G) Quantification of sections as represented in part F. (n = 5-7/group). RT-qPCR comparisons in adipogenic (H) and Wnt-related (I) genes from the secondary spongiosa of femurs as shown in F (n = 3-5/group). Statistics analyzed by 2-tailed student t-test, and lines plotted reflect the mean ± SEM, * p<0.05, ** p<0.005, *** p<0.0005

Bone marrow stem/stromal cells (BMSCs) represent common precursors for adipocytes and osteoblasts, and marrow adipose tissue (MAT) volume is known to be inversely correlated with trabecular bone mass (Ko et al., 2021; Pierce et al., 2019; Tencerova et al., 2018; Yue et al., 2016). To determine if MAT volume is affected in the *Ksr2* KO mice, we evaluated the levels of osmium tetroxide retaining MAT in femurs of 28-week-old mice by microCT (Figure 4D). MAT volume was significantly reduced in all three compartments (proximal and distal metaphysis, and diaphysis) in the tibia of *Ksr2* KO mice compared with controls (Figure 4E). Consistent with these data, there was a significant reduction in adipocytes in the distal femoral metaphysis of *Ksr2* KO mice (Figure 4, F and G).

To determine the potential regulatory molecules that contribute to changes in MAT in *Ksr2* KO mice, we compared mRNA expression of adipocyte markers at the trabecular compartment in distal femurs of *Ksr2* KO and control mice. Neither markers associated with white adipocytes (*Fabp4*, *Slc7a10*), or brown adipocytes (*Ucp1*, *Prdm16*) were different in *Ksr2* KO femurs, while of the adipogenic regulatory transcription factors evaluated, *Pparg* was mildly but significantly reduced, while *Cebpa* was not changed. However, key adipokines (*Lep, Cfd*) were decreased (Figure 4H). Since Wnt signaling is critically involved in regulating adipocyte differentiation, we also measured mRNA levels of several Wnt-related genes, but only found a decrease in *Wnt8b*, and an increase in *Ccnd1*, a Wnt target gene, in the bones of *Ksr2* KO mice (Figure 4I). Thus, changes in adipokine gene expression in both body and MAT adipocytes are altered when *Ksr2* is deleted globally.

### Loss of Ksr2 delays femoral fracture healing and results in more fragile bones

The pathological obese/T2D condition predisposes bones to compromised fracture healing and increased fracture risks. Since the absence of *Ksr2* results in increased appendicular bone deposition, we evaluated whether this increased rate of bone formation would prove beneficial in *Ksr2* KO mice. Healing response was compared between WT and KO mice at 16 weeks of age following stabilized closed femoral fractures (Figure 5A). X-ray analysis 3 weeks after fracture showed improved bony union of the callus in WT mice and increased callus size in KO mice (Figure 5B).

**Figure 5.**
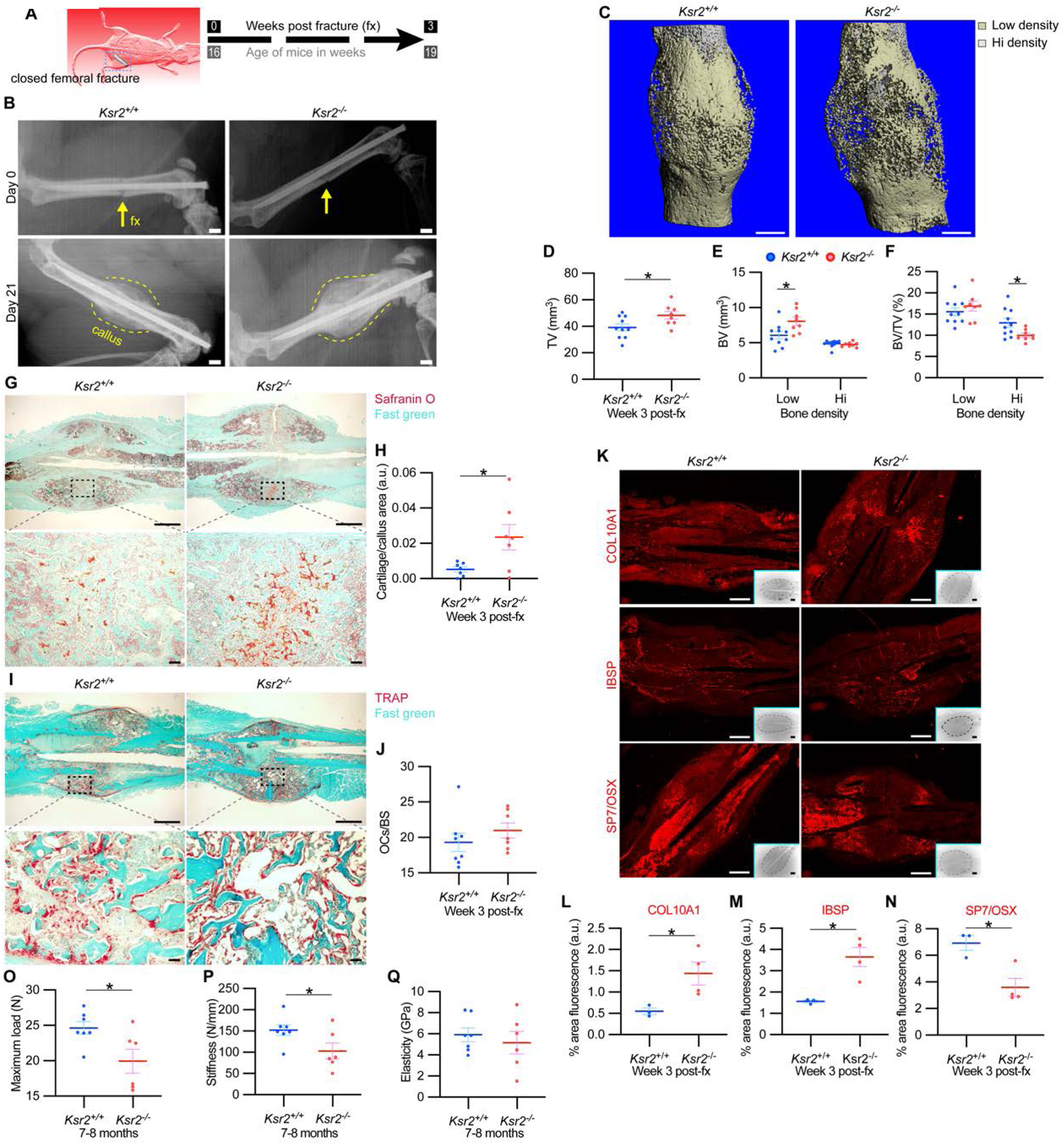
Delayed fracture healing but increased fragility in *Ksr2* knockout mice. (A) Schematic of strategy. (B) Representative X-ray images of bones that underwent closed mid-femoral fracture in WT and *Ksr2^-/-^* KO mice on the day of surgery (day 0) and day 21 post-fracture (fx). Yellow arrows point to induced fracture, while calluses are outlined by dotted yellow lines. Scale bar: 1mm. (C) Representative 3D-microCT reconstruction images of fracture calluses at 3 weeks post-fx. Color-coded differences in bone density are indicated in the legend. (D-F) Quantification of microCT data for TV, total volume; BV, bone volume, and BV/TV (n = 8-10/group). (G) Representative Safranin O stained chondrocytes in WT and *Ksr2^-/-^* bones at 3 weeks post-fx showing increased cartilage in KO mice, and corresponding quantification (H), (n = 6-7/group). (I) Representative TRAP-stained osteoclasts in calluses at 3 weeks post-fx., showing no difference between genotypes, with corresponding quantification of osteoclasts/bone surface within callus (H) (n = 7-8/group). All histology sections were counterstained with fast green dye. Scale bars: 1mm at low magnification (mag)- top rows, or 100μm at high mag-bottom rows. (K) Representative immunofluorescence images for COL10A1, IBSP, SP7/OSX of fracture callus at 3 weeks post-fx. Scale bar: 1mm at low mag insets; 100μm at high mag. (L-N) Quantitation of fracture callus. (O-Q) Three-point bending test show femurs of *Ksr2*^-/-^ KO mice tolerate less load to fracture with reduced stiffness, while elasticity remains unchanged. N, Newton; GPa, GigaPascal. (n=6-7/group mixed genders; 2 males per group). Statistics were analyzed by 2-tailed student t-test, and graphed lines represent the mean ± SEM, * p<0.05.

MicroCT measurements also showed increased total volume in the fracture callus of KO mice (Figure 5, C and D). When the callus was segmented into low and high densities for analysis, an increase in low density woven bone volume was observed in *Ksr2* KO fracture callus but no changes were observed in high density cortical bone volume (Figure 5, E and F). Consequently, the low density callus BV/TV in the *Ksr2* KO fractures was not significantly different from WT, while the high density BV/TV callus was reduced in *Ksr2* KO mice (Figure 5, E and F). Therefore, KO mice exhibited a larger fracture callus with increased low density woven bone and reduced high density cortical bone, consistent with delayed fracture callus development.

Histological evaluation also revealed increased Safranin-O-stained cartilage in *Ksr2* KO mice but no difference in TRAP+ osteoclasts compared to WT mice (Figure 5G-J), suggesting that the differences observed between WT and KO were not caused by differences in bone resorption, but rather by delayed endochondral ossification. Proteins associated with hypertrophic chondrocytes, COL10A1 and IBSP (Gomez et al., 2022), were also increased in *Ksr2* KO mice, while SP7 was reduced, indicating that the delay in bone formation occurred during the ossification of the hypertrophic cartilage (Figure 5K-N). Combined, these results show that although deletion of *Ksr2* leads to obesity with increased bone mass, fracture healing is compromised despite the increased bone accretion observed in unfractured bones.

Since increased trabecular bone mass associated with obesity/T2D in humans is paradoxically associated with an increased risk of fracture (Greco et al., 2015; C. Ma et al., 2018; Moseley, 2012; Oei et al., 2013), femoral bones of WT and KO mice were tested for resistance to fracture by a three-point bending test. A lighter load was required to break KO bones, which displayed reduced stiffness, yet no change in elasticity (Figure 5O-Q). Therefore, as with humans, the femoral bones of *Ksr2* KO mice likely have structural integrity deficits that are more prone to fracture.

### Ksr2 is sufficient to inhibit osteoblast but not osteoclast differentiation

While *Ksr2* was known to be highly expressed in the brain, its expression in bone was unknown. By immunofluorescence using longitudinal sections of distal femoral bone sections from 3-week-old WT mice, positive KSR2 staining was observed at the epiphyseal secondary ossification center, and the metaphyseal region encompassing the primary and secondary spongiosa, coinciding with SPP1 (Figure 6A). To further explore if *Ksr2* is expressed in osteoblasts and/or osteoclasts, we evaluated whether *Ksr2* mRNA is expressed during osteoblast or osteoclast differentiation from primary pre-osteoblasts or macrophages isolated from WT calvarial or femoral bones, respectively. The fidelity of differentiation in each condition was reflected by temporal upregulation of *Alpl* mRNA in osteoblasts or *Acp5* mRNA in osteoclasts, relative to vehicle-treated controls (Figure 6, B and C). In osteoblasts, *Ksr2* exhibits a biphasic response, being inhibited on day 3 and transiently upregulated on day 14, before returning to basal levels on day 21, while *Ksr1* levels hovered around the baseline (Figure 6B). By comparison, *Ksr2* was only upregulated at the end of differentiation in osteoclasts (Figure 6C). Combined, this data shows that KSR2 is expressed in osteoblasts in vivo, and during osteoblast and osteoclast differentiation ex vivo.

**Figure 6.**
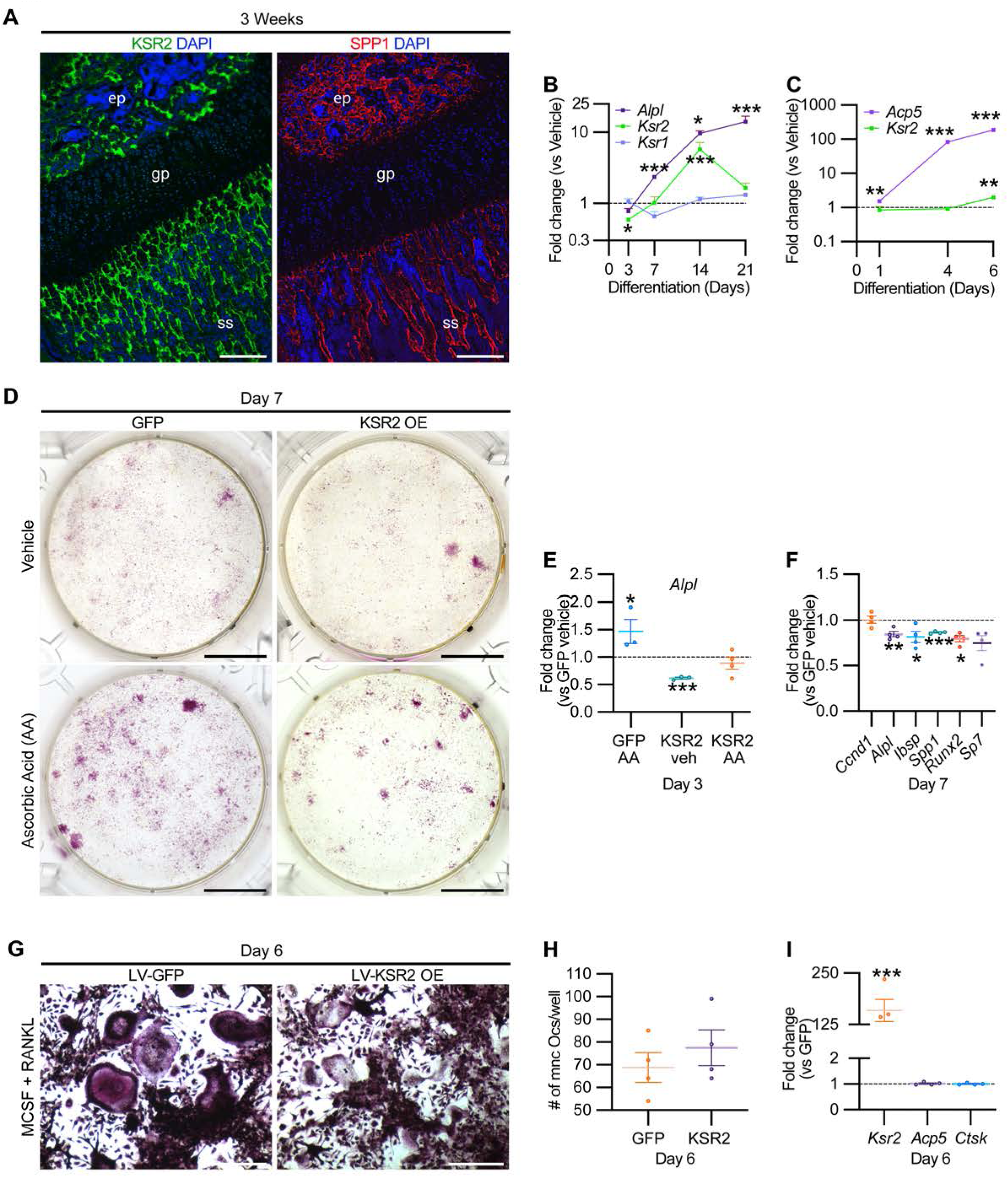
KSR2 is expressed in bone, and ex vivo gain of function studies demonstrate *Ksr2* represses osteoblast differentiation but is dispensable for osteoclast differentiation. (A) Representative immunofluorescence for KSR2 (green) and OPN (red) expression in longitudinal sections of distal femur epiphysis from 3-week-old WT mice, counterstained with DAPI (blue). Abbreviations: ep, epiphyseal bone; gp, growth plate; ss, secondary spongiosa. Scale bar: 100μm. (B) Ex vivo time-course RT-qPCR characterization of *Alpl*, *Ksr2*, and *Ksr1* mRNA expression on calvaria pre-osteoblasts isolated from WT mice following induction with osteoblast differentiation conditions relative to vehicle treatment (n = 3-4 independent experiments). (C) Ex vivo time-course RT-qPCR characterization of *Acp5* and *Ksr2* on primary macrophage cells isolated from femoral bones of WT mice following osteoclast differentiation relative to vehicle treatment. (n = 3-4 independent experiments). (D) Representative images of Alizarin red-stained primary bone marrow stromal cells with forced expression of either GFP or KSR2 after 7 days of treatment with either Vehicle or Ascorbic Acid (AA). Scale bar: 10mm. (E) RT-qPCR for *Alpl* on day 3 or (F) various osteoblast markers on day 7 (n = 4 independent experiments). (G) Representative images of multinuclear osteoclasts stained with ACP5/TRAP following 6 days of osteoclast differentiation from macrophages isolated from femurs and transduced with either GFP or KSR2. Scale bar: 100μm (H) Quantification of multinuclear osteoclasts (Ocs) counted/well as shown in part G. (I) RT-qPCR for *Ksr2*, or osteoclast markers *Acp5*, *Ctsk* in osteoclasts on day 6 of differentiation as represented in part G. Statistics analyzed by 2-tailed student t-test, and graphed lines represent the mean ± SEM, * p<0.05, ** p<0.005, *** p<0.0005

Therefore, we assessed whether Ksr2 has any effect on the differentiation of either of these two lineages by a gain of function approach. BMSCs or macrophages isolated from femurs of 3-week-old WT mice were transduced with lentiviral vectors encoding either the KSR2 open reading frame, or GFP as controls, and evaluated for differences in differentiation potential. BMSCs harboring GFP or overexpressed (OE) KSR2 (10-30-fold), underwent osteoblast differentiation in the presence of ascorbic acid (AA) or vehicle for 7 days. Alizarin red staining shows reduced differentiation in KSR2 OE cultures compared to GFP controls (Figure 6D). Ksr2 OE also reduced the expression of osteoblast differentiation markers *Alpl*, *Ibsp*, and *Spp1*, as well as *Runx2* and *Sp7,* while *Ccnd1* was not changed (Figure 6, E and F), suggesting KSR2 exerts direct effects on osteoblast differentiation.

Macrophage differentiation toward the osteoclast lineage was achieved in both GFP and KSR2 OE cells as noted by the presence of TRAP-stained multinuclear osteoclasts in both populations on day 6 (Figure 6G), and their quantification resulted in minimal differences (Figure 6H). Also, we found no changes in *Acp5* or *Ctsk* between these cells, while *Ksr2* continued to be overexpressed (Figure 6I). Thus, this gain of function strategy suggests KSR2 negatively regulates osteoblast differentiation from BMSCs but is likely dispensable for osteoclast differentiation.

### Ksr2 regulates bone formation autonomously

To formally address whether central hypothalamic KSR2 mediated obesity might also regulate distal limb bone formation non-autonomously, we took a two-pronged approach. In the first approach, *Ksr2* KO mice were split into two groups, one was allowed to feed ad libitum (Ad lib), while the other group was pair-fed according to amounts eaten by WT mice for 12 consecutive weeks starting at 4 weeks after birth. As reported previously (Revelli et al.), *Ksr2* KO mice consumed twice as much food on average compared to WT mice. Also, serum leptin levels were increased several-fold in the Ksr2 KO mice, which was rescued by pair-feeding according to amount of food eaten by WT control mice (Revelli et al., 2011). However, serum leptin levels were not measured in this study. Ad lib-fed KO mice again showed significant gains in BW, percent body fat, and femoral BMD relative to WTs, while pair-fed KO mice neither gained weight nor body fat but retained the increased femoral BMD (Figure 7A-C). Consistent with these data, trabecular bone volume fraction measured by microCT was significantly higher in *Ksr2* KO mice than WT controls after pair feeding (Figure 7D).

**Figure 7.**
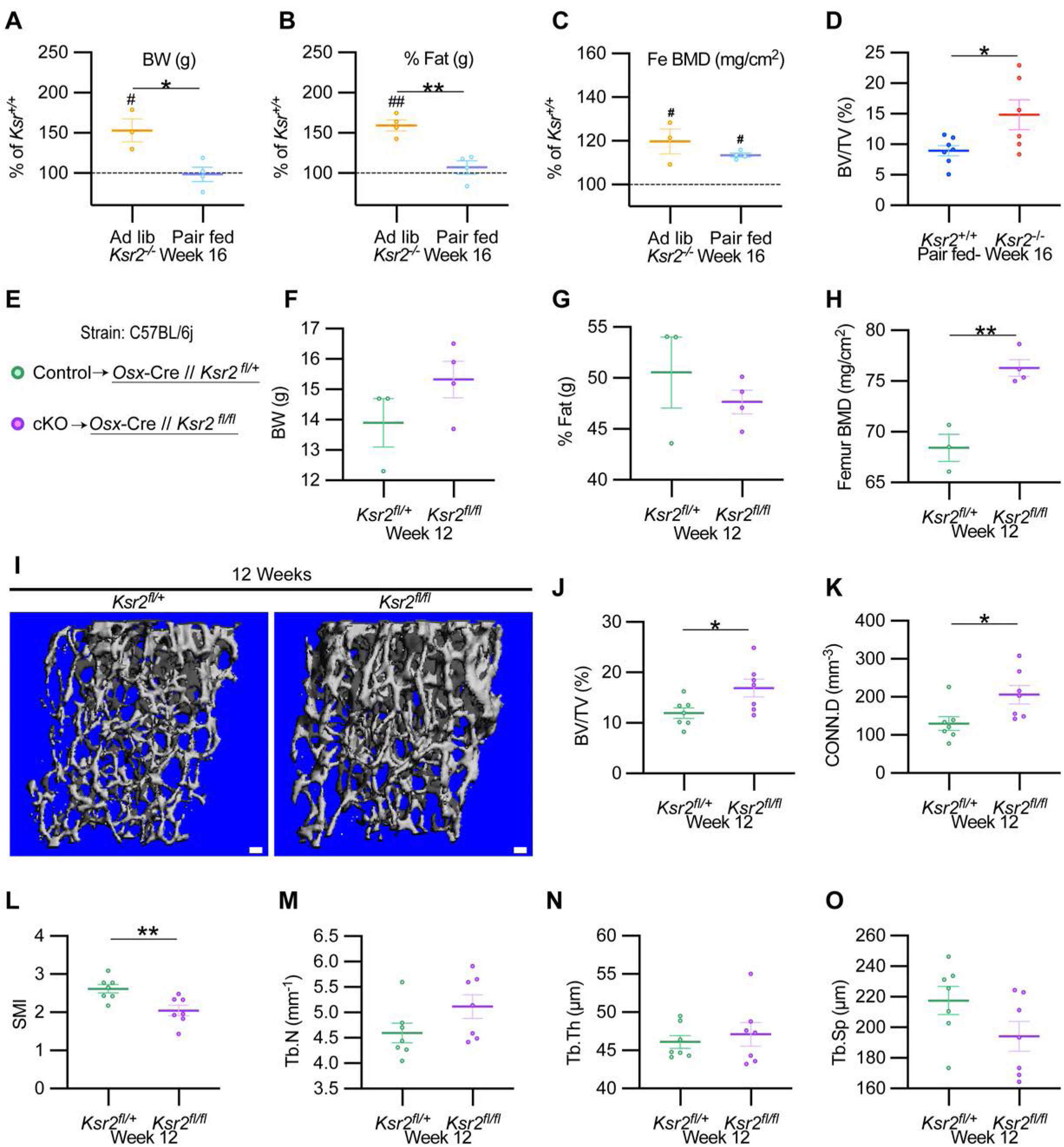
*Ksr2* regulates femoral trabecular bone autonomously. (A-C) Pair feeding experiments reveal that gains in mineral density are acquired independently of eating-induced weight gains in KO mice fed at will (Ad lib) or pair-fed according to the amount eaten by WT mice. Parts A-C are represented as a percentage relative to WT. Abbreviations: BW, Body weight; Fe BMD, femur bone mineral density; BV/TV, Bone volume/total volume. (n = 3-6/group). (D) BV/TV from femoral metaphysis of WT and KO at end of pair feeding. (E) Conditional knockout strategy. (F-H) Differences between Control (*Ksr2^fl/+^*) and conditional knockout, cKO (*Ksr2^fl/fl^*), mice in percent body fat (F), body weight (G), and femur bone mineral density (H) (n = 3-4/group). (Note: B,C,G,H reflect dual-energy x-ray absorptiometry measurements). (I) Representative 3D microCT reconstruction images of distal femoral metaphysis in Control and cKO mice at 12 weeks of age, revealing increased trabecular bone in cKO mice. Scale bar: 100μm. (J-O) MicroCT measurements from the trabecular bone as represented in part I (n = 7 mice per group; mixed genders). Abbreviations: BV/TV, Bone Volume/Tissue Volume; CONN.D, Connectivity Density; SMI, Structural Model Index; Tb.N, Trabecular number; Tb.Th, Trabecular thickness; Tb.Sp, Trabecular spacing. Statistics were analyzed by 2-tailed student t-test, and graphed lines represent the mean ± SEM, * p<0.05, ** p<0.005 for comparisons between groups labeled on the x-axis. In parts A-C, significance between *Ksr2* KO and WT for a given condition is represented by # p<0.05, or ## p<0.005.

In the second approach, *Ksr2* was conditionally deleted in osteoblasts via *Sp7/Osterix-Cre* mice which have been successfully used for disrupting gene function in osteoblast lineage cells (Buettmann et al., 2019; Ko et al., 2021). Since the *Osx-Cre* transgenic mice exhibit a mild skeletal phenotype (W. Huang and Olsen, 2015), we used *Osx*-*Cre^+^ Ksr2* floxed heterozygous mice as controls (Figure 7E). While neither body weight nor percent fat was different between *Osx*-*Cre^+^ Ksr2* floxed heterozygous (control) and homozygous (conditional KO, cKO) mice, femoral BMD was significantly increased in the *Ksr2* cKO mice compared to control mice (Figure 7F-H). In sync, microCT analyses of distal femoral metaphyseal secondary spongiosa of cKO mice exhibited similar osteal gains relative to controls, as observed between global KO and WT mice (Figure 7I-O). Therefore, this data indicates that gains in bone mass can be regulated autonomously by KSR2 expressed in bone, independent of the centrally regulated effects of *Ksr2* in the hypothalamus.

### Ksr2 affects osteoblast differentiation through mTOR signaling

Mechanistically, *Ksr2* has been shown to regulate changes in visceral fat by multiple mechanisms in the hypothalamus including AMPK and mTOR signaling (Figure 8I) (Costanzo-Garvey et al., 2009; Pearce et al., 2013; Revelli et al., 2011). In the ST2 mouse stromal cell line, KSR2 OE inhibited *Alpl* expression in both normal and high glucose media as well as in the presence or absence of insulin treatment (Figure 8A).

**Figure 8.**
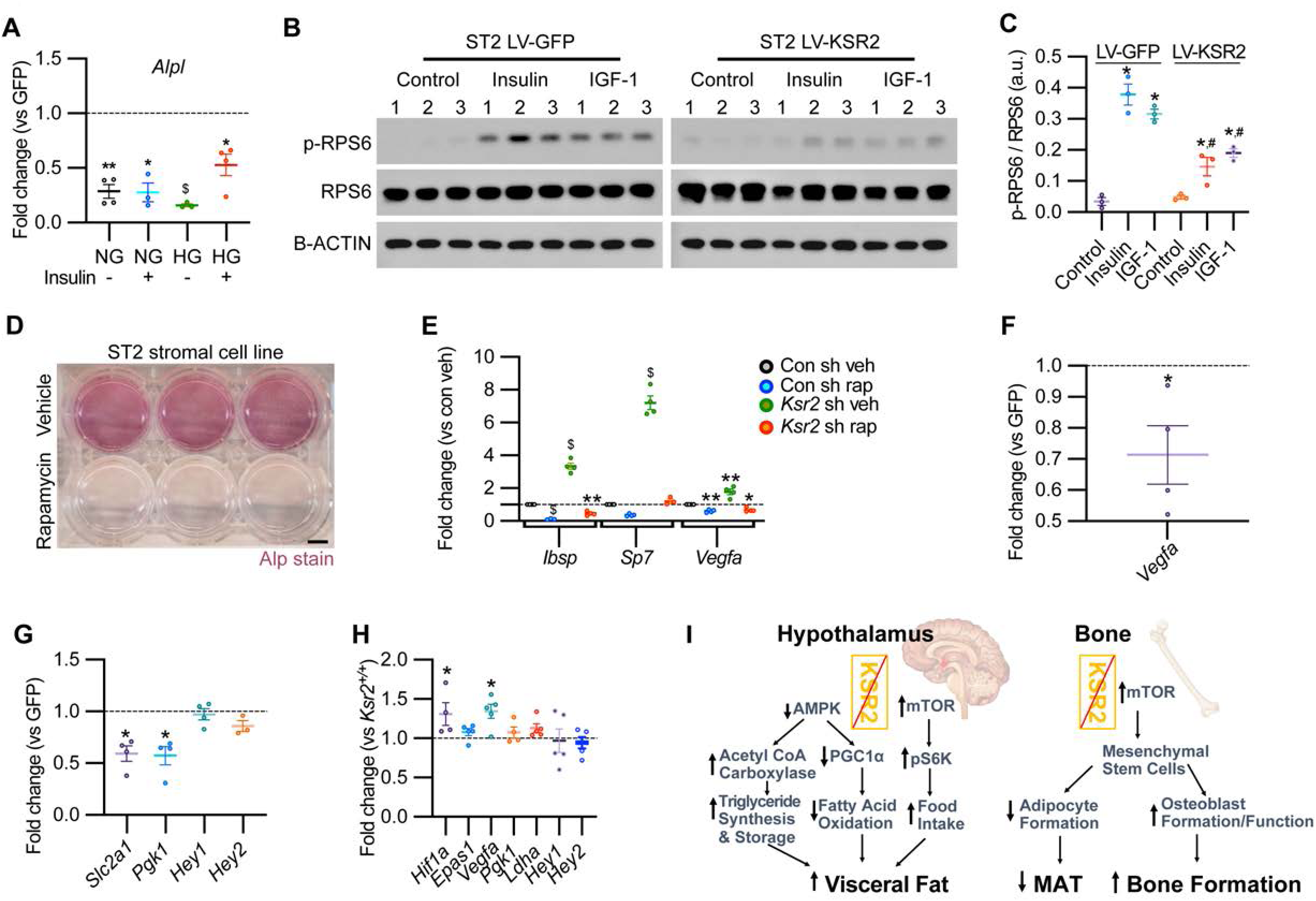
KSR2 promotes osteoblast differentiation through mTOR signaling affecting *Hif1a* and *Vegfa,* but not *Notch* signaling in the process. (A) RT-qPCR for *Alp* from ST2 stromal cells overexpressing KSR2 or GFP by Lentivirus (LV) on day 3 in normal glucose (NG) or high glucose (HG) without (-) or with (+) insulin. (B) Western blot for ST2 cells overexpressing GFP or KSR2 after 30 minutes in vehicle Control, 100 μg/ml Insulin, or 100ng/ml IGF-1, for the mTOR response target phosphorylated p-RPS6, total RPS6, or loading control, ∝-ACTIN; 1,2,3 indicate biological replicates. (C) Quantification of WB comparing p-RPS6/RPS6 ratios. Within group (*), between groups (#) comparisons (n = 3/group). (D) Representative image of ALP activity for ST2 stromal cells in osteoblast differentiation conditions on day 7 treated with vehicle (top row) or 10nM Rapamycin (bottom row). Scale bar: 10mm. (E) RT-qPCR quantification of ST2 cells transduced with empty vector Control (Con) or *Ksr2* shRNA and treated with either vehicle or 10nM Rapamycin following 48 hours of osteoblast differentiation. (n = 4/group). (F, G) RT-qPCR from ST2 stromal cells with KSR2 overexpression following 72 hours of osteoblast differentiation, plotted as a function of level detected in GFP controls. (n = 4/group). (H) RT-qPCR on genes related with Hypoxia or Notch signaling on RNA extracted from whole femurs of 12-week-old WT or *Ksr2* KO mice. Values represent fold change for KO relative to WT (set to 1, dashed line). (I) Model diagram summarizing results where high levels of KSR2 lead to low levels of mTOR activity resulting in low bone density, while the absence of KSR2 results in high levels of mTOR activity resulting in high bone density. All statistics analyzed by 2-tailed student t-test, graphed lines represent mean ± SEM. # p<0.05, * p<0.05, ** p<0.005, $ p<10^-6^

In ST2 cells transduced with GFP control vector, both Insulin and IGF-1 promoted a marked increase in the activated form of phosphorylated (pRPS6) relative to total RPS6 compared with vehicle-treated control cells, as expected. By contrast, pRPS6 activation was significantly reduced in KSR2 OE ST2 cells (Figure 8, B and C). AMPK phosphorylation was unaffected by *Ksr2* OE in ST2 cells (data not shown). mTOR is known to promote osteoblast differentiation in ST2 cells (Chen et al., 2014), and treatment of these cells under osteoblast differentiation conditions with the classic mTOR inhibitor, rapamycin, significantly blocked differentiation measured by ALPL staining (Figure 8D). To further determine whether *Ksr2* signals through mTOR in stromal cells during osteoblast differentiation, ST2 cells were knocked down with *Ksr2* shRNA or a non-specific control shRNA, and osteoblast differentiation was tested with either rapamycin or vehicle control and evaluated after 48 hours for expression of *Ibsp*, *Sp7* or *Vegfa* by RT-qPCR. Increased expression of bone formation markers by lentiviral *Ksr2* shRNA treated cultures is abolished by rapamycin treatment (Figure 8E). These results indicate that *Ksr2* regulates osteoblast differentiation via mTOR activation.

To determine the downstream targets of KSR2/mTOR, we measured expression levels of Notch and hypoxia signaling genes (Bjedov and Rallis, 2020; Frey et al., 2014; B. Huang et al., 2015) in ST2 cells overexpressing KSR2 or GFP. We found that Ksr2 OE reduced the expression of hypoxia signaling targets (*Vegfa, Slc2a1*, *Pgk1*) but did not affect Notch targets (*Hey1*, *Hey2*), (Figure 8, F and G). Accordingly, crucial hypoxia markers (*Hif1a*, *Vegfa*) were increased in the bones of 12-week-old *Ksr2* KO mice, but Notch targets were not changed (Figure 8H).

## Discussion

The impact of obesity on bone health is an area of significant concern. While obesity is known to exert complex effects on bone mass and skeletal fragility, the mechanisms by which obesity influences bone metabolism are not well understood. In this study, we used a *Ksr2* KO genetic mouse model to investigate the relationship between obesity and bone health. We found that the distal femoral metaphyseal trabecular bone was considerably denser in *Ksr2* KO compared to littermate WT controls in two genetic backgrounds, while deletion of *Ksr1* did not affect trabecular bone mass. At the cellular level, we found by histomorphometric analysis, *ex vivo*, and *in vitro* studies via *Ksr2* gain and loss of function that KSR2 is a negative regulator of trabecular bone formation that may act by controlling differentiation of mesenchymal stem cells to the osteoblast or adipocyte fate. Although analysis of serum markers did not concur, this was likely because serum was evaluated at the advanced age of 26 weeks, when the skeletal phenotype is already manifested in the *Ksr2* KO mice. By contrast, KSR2 did not significantly affect osteoclast formation or functions. Moreover, our data from *Ksr2* KO mice that were pair-fed similar to WT mice and osteoblast-specific *Ksr2* conditional KO mice revealed that *Ksr2* controls osteoblast differentiation autonomously. At the molecular level, our findings demonstrate that *Ksr2* negatively regulates osteoblast differentiation by repressing mTOR activity (Figure 8I). Since global *Ksr2* KO mice exhibited delayed fracture healing with an arrest at endochondral ossification, as observed in obese humans with T2D, and since *Ksr2* genetic polymorphisms are linked to obesity, an understanding of the cellular and molecular pathways of KSR2 regulation of mesenchymal stem cell fate may have important ramifications in promoting bone health in obese individuals.

Genetic differences between mouse strains are known to affect their susceptibility to gain excess weight, with the C57BL/6J strain being more vulnerable, and therefore used more often than DBA/1LacJ to study diet-induced weight gain (Garofalo et al., 2003; Linder, 2006). Also, the impact of genetic background on gene deletion phenotypes is well established (Doetschman, 2009; Linder, 2006). In accord, we noticed that *Ksr2* KO mice in the DBA background lived longer than those in the C57 background (data not shown). Since *Ksr2* deletion results in weight gain and increased femoral bone mass in both strains, it is highly likely that *Ksr2* also regulates bone formation in other genetic backgrounds. This warrants further investigation in other mammals and, in particular, in humans. Furthermore, our data show that disruption of the *Ksr2* gene had no significant effect on trabecular bone mass in the vertebra. These data are consistent with what is known in the literature that the heritability of bone mineral density varies across skeletal sites (Kemp et al., 2014; Rowe et al., 2018) and differences in mechanisms that regulate bone accretion in long bones versus vertebrae.

Other monogenetic models that lead to obesity/T2D defects have also investigated bone phenotypes. In particular, knockouts of different genes in the leptin-melanocortin feedback loop that signals satiety in the hypothalamus, generally result in obesity, but although the neural circuits are unidirectional, both anabolic and catabolic effects have been observed in bone. For instance, while leptin KO mice result in reduced femur length and BMD (Steppan et al., 2000; X. Wang et al., 2007), they have increased vertebral bone mass (Ducy et al., 2000), consistent with the idea that genes can have distinct effects in different anatomical regions. By contrast, *Mc4r* and *Npy1r* knockouts result in increased femoral BMD (Ahn et al., 2006; Baldock et al., 2007; Braun et al., 2012), while *Mc3r* knockouts have reduced femur length and BMD (Lee et al., 2016). Many of these genes, such as *Npy1R* and *Mc4r* are expressed in both hypothalamic neurons and osteoblasts (Baldock et al., 2007; Q. Zhong et al., 2005), which may be partly responsible for the complex skeletal phenotypes seen in these mice. Similarly, *Ksr2* is expressed in the hypothalamus, and in this study, we found that it is also expressed and functional in cells of the osteoblastic lineage. Although our pair-feeding and conditional knockout studies define a role for KSR2 function in bone that can be dissociated from its hypothalamic function, *Ksr2* global knockout and likely humans with *Ksr2* genetic polymorphisms have malfunctions of both hypothalamic KSR2 regulated food intake as well as bone KSR2 regulated osteoblast formation. In future studies, we will address whether *Ksr2* plays similar or different roles in other skeletal sites and whether conditional hypothalamic deletion of *Ksr2* has any effect on bone physiology. An alternative means to study the effect of obesity on bone is provided by diet-induced obesity models. Generally, these mice result in excess body fat, with reduced trabecular bone mass at the expense of increased MAT (Bonnet et al., 2014; Scheller et al., 2016; Tencerova et al., 2018). This would suggest that excess body adipocytes, which secrete adipokines that are known to influence different aspects of bone maintenance by regulating the differentiation or function of BMSCs, osteoblasts, or osteoclasts. Consistent with the idea that adipocyte-derived factors regulate osteoblast and osteoclast functions are the findings that bone mass is increased under conditions of generalized reduction in adipose tissue, as in the case of congenital lipodystrophy (Zou et al., 2019). However, the local secretion of adiponectin, an adipokine, in MAT appears to provide a stronger influence in this model (L. Zhong et al., 2020). Consistently, we did detect a decrease in adiponectin expression in the femurs of *Ksr2* KO mice (data not shown). Nevertheless, this indicates that adipocytes can influence bone homeostasis by both systemic and local signals.

Leptin is another well-known adipokine secreted by adipocytes, that is increased in obese animals and is known to regulate bone formation. However, a significant role for adipocyte-derived leptin in mediating the gains in trabecular bone mass in *Ksr2* KO mice does not seem likely. We previously found that Leptin is increased in serum of *Ksr2* KO mice, but Leptin resistance was not causative of weight gains in these nice (Costanzo-Garvey et al., 2009; Revelli et al., 2011). Here we show that *leptin* mRNA is increased in body fat depots but believe this is also not responsible for increased bone mass. Our data shows that pair-fed *Ksr2* KO mice and osteoblast-specific conditional *Ksr2* KO mice do not gain excess adipocytes, yet still resulted in gains in trabecular bone. Since leptin is produced by fat, which did not change in either of these conditions, these results suggest gains in systemic leptin produced by excess fat may not be a causative factor in bone mass accretion when *Ksr2* is deleted. Interestingly, while we did see a reduction in *leptin* mRNA expression in femurs, in isolation this might be expected to reduce bone mass, given that *in vitro* and *in vivo* reports indicate that leptin is anabolic to limb bones (Astudillo et al., 2008; Gordeladze et al., 2002; Steppan et al., 2000; X. Wang et al., 2007). While it has been reported that *leptin* mRNA is higher in visceral adipose tissue than BMAT (Liu et al., 2011), in the absence of KSR2, it is reduced even further, yet this does not result in bone loss.

Interestingly, MAT is significantly reduced in the long bones of *Ksr2* KO mice compared to WT mice as revealed by Osmium tetroxide micro-CT evaluation, as well as histological analyses of adipocyte numbers. By contrast, MAT has been shown to be increased in mice with disruption of leptin, leptin receptor, as well as mice fed with high-fat diets, all three models that show reduced femoral trabecular bone mass (Hamrick et al., 2004; Tencerova et al., 2018; Yue et al., 2016). These findings together with the known fact that mesenchymal stem cells represent common precursors of both osteoblasts and adipocytes raise the possibility that KSR2 might modulate the switch between osteoblast and adipocyte differentiation produced by mesenchymal stem cells, and, thereby, bone formation and MAT. Further studies are required to determine the cause-and-effect relationship between changes in *Ksr2* expression and regulation of mesenchymal stem cell differentiation. In this regard, a recent study demonstrated that complement factor D/adipsin from bone marrow adipocytes regulates bone marrow stromal cell fate determination through activation of the complement system (Aaron et al., 2021). We, therefore, examined if the expression of adipsin was altered in the bones of *Ksr2* KO mice and found reduced expression of adipsin in both body fat and femoral adipocytes. The issue of whether KSR2 regulates adipsin expression directly or indirectly via other factors remains to be established. While Aaron et al. demonstrated that adipsin is a downstream target of PPARG, *Pparg* transcription was only mildly reduced in *Ksr2* KO bones, thus raising the possibility that KSR2 might regulate adipsin expression independently of PPARG.

Other factors that might affect bone physiology in obese and T2D conditions are inflammatory cytokines. While adipocytes are known to contribute to increased levels of pro-inflammatory cytokines such as TNF, Il6, Il17, and Tnfsf11/RANKL (Benova and Tencerova, 2020; Kawai et al., 2021) during certain pathological states that can promote increased osteoclastogenesis and resorptive activity, we did not see changes in bone resorption in *Ksr2* KO mice. However, increased levels of these inflammatory cytokines could be responsible for the altered fracture healing in *Ksr2* KO mice. Diet-induced obesity models have reported reduced callus bone volume and increased marrow adiposity, possibly due to a faster rate of callus resorption (Brown et al., 2014), producing bones with microstructural deficits in collagen matrix and increased advanced glycation end products (Khajuria et al., 2020). Although some studies find smaller callus in obesity/T2D fracture callus (Brown et al., 2014), others have also observed increased callus size in the fracture callus of DIO-obesity/T2D model fractures, with increased hypertrophic chondrocytes, and delayed fracture healing, similar to the results reported here (Marin et al., 2021). It is possible that fractures using an intramedullary pin is not as stabilized in the *Ksr2* KO mice as that of WTs because of increased body weight, thus leading to a larger less dense callus, a phenomenon frequently seen in non-stabilized human fractures. Further time-course studies are needed to determine the cause for the delayed remodeling of fracture callus in the *Ksr2* KO mice, and if the healed bone in *Ksr2* KO mice are mechanically weaker than the healed bones of control mice. Regardless, the diabetic state results in a deranged inflammatory condition that is believed to affect the vascular system by the production of advanced glycation end products and multiple factors that may delay fracture healing (Marin et al., 2018), and we show here that loss of *Ksr2* may not be sufficient to impart improved fracture healing for diabetics. In the future, we will determine if increased general adiposity is the cause of delayed fracture healing in global *Ksr2* KO mice by evaluating if fracture healing is affected in mice with conditional disruption of *Ksr2* in osteoblasts. Nevertheless, bones of *Ksr2* KO mice were less resistant to fracture, in agreement with the observation in obese/T2D humans which present with increased bone mass and fracture susceptibility (Greco et al., 2015; C. Ma et al., 2018; Moseley, 2012; Oei et al., 2013).

mTOR regulation of osteoblast formation remains controversial, as both positive and negative associations have been reported (Chen et al., 2014; Martin et al., 2010; Martin et al., 2015; Xian et al., 2012; Yeo et al., 2021). Our in vitro results show that forced expression of KSR2 reduces mTOR signaling, while knockdown of *Ksr2* promotes induction of osteogenic factors, but not when mTOR signaling is inhibited by rapamycin. Moreover, both in vitro and in vivo results suggest KSR2 and mTOR affect hypoxia but not Notch signaling genes. The anabolic effects of hypoxia signaling on bone mass and vasculature is well established (Mohan and Kesavan, 2022; Shen et al., 2009; Wan et al., 2010; Y. Wang et al., 2007; Wolf et al., 2022). Thus, whether the increased hypoxia signaling pathway observed in bones of *Ksr2* KO mice contributes to the increased trabecular bone mass remains to be established. Thus, this work implicates mTOR as a positive effector of osteoblast differentiation, that can be regulated by KSR2 (Figure 8I). Future studies are needed to determine how KSR2 regulates mTOR signaling biochemically, whether KSR2 regulates BMSC fate decision via mTOR in either the mTORC1 or mTORC2 complex, which is reportedly one means of affecting BMSC fate regulation (Martin et al., 2015; Sen et al., 2014), and whether osteoblast-specific deletion of mTOR in *Ksr2* KO mice will reverse the bone gains in *Ksr2* KO mice.

In summary, our investigation of bones in *Ksr2* knockout genetic mouse models resulted in the identification of a novel animal model in which the obesity/T2D condition coincides with increased appendicular bone mass. Since *Ksr2* genetic polymorphisms are linked to obesity/T2D in humans, our full understanding of how KSR2 differentially regulates general tissue adiposity versus bone marrow adiposity could lead to the identification of novel therapeutic strategies to promote bone health in humans with obesity/T2D.

## Methods

### Mice

Femoral bones of mice in the C57BL/6J-Tyr*^c-Brd^* X 129^SvEvBrd^ hybrid background were transferred from Lexicon pharmaceuticals to the Veteran’s Affairs Loma Linda Healthcare System (VALLHS) and analyzed at VALLHS. *Ksr2*^+/-^ mice in the DBA/1LacJ were transferred from the University of Nebraska to the VALLHS and maintained by inbreeding for experimentation and further analysis. *Ksr2*-floxed mice were generated by insertion of LoxP sites flanking exon3 of *Ksr2* as described (Guo et al., 2017), and mated to *Sp7/Osx-*Cre mice (a kind gift from Dr. Andrew P. McMahon, University of Southern California, USA) for osteoblast-specific deletion of *Ksr2*. *Ksr1*^+/-^ mice were a kind gift from Dr. Andrey S. Shaw at Washington University School of Medicine (St. Louis, USA), and were bred to purity in the C57BL/6J background for bone analysis. Mice genotyping was done by conventional tail snip PCR with DNA primers. All animals were housed at the animal facility of VALLHS (Loma Linda, CA, USA) according to approved standards with controlled temperature (22°C) and illumination (14-hour light, 10-hour dark). Mice were fed a standard chow diet. The approved anesthetic (isoflurane) was used for anesthesia, and CO_2_ exposure was used for euthanasia followed by cervical dislocation.

### Pair feeding

Pair feeding studies were performed as described (Pearce et al., 2013; Revelli et al., 2011). Mice were fed a standard chow diet throughout the experiment.

### Fractures

At 16 weeks of age, *Ksr2* KO and WT mice of mixed genders were subjected to stabilized closed femoral fracture by a modification of the three-point bending approach (Rundle et al., 2008). Fracture tissues were harvested at 3 weeks post-fracture for further analysis when bony callus union is expected in this model and after which fracture callus remodeling should normally complete healing.

### microCT

Femur lengths, trabecular and cortical bone parameters were measured on a VIVA CT40 (Scanco medical, Bruttisellen, Switzerland) micro-computed tomography (microCT) system. Bones were fixed in 10% formalin overnight, washed, and imaged in 1X PBS with 55 to 70 kVp volts at a voxel size of 10.5μm. Images were reconstructed using the 2D and 3D image software provided by the Scanco VIVA-CT 40 instrument (Scanco USA, Wayne, PA). For analysis of the spine, bones were sampled at the 4^th^ Lumbar (L4) vertebrae. Osmium tetroxide experiments were performed for the measurement of marrow adipose tissue as described (Lindsey, Godwin, et al., 2019).

### Dual X-ray absroptiometry

Total body bone mineral density, percent body fat, femoral bone mineral density, and X-ray fracture images were analyzed on a Faxitron Radiography system (Hologic, Bedford, MA). Images were acquired with 20 kV X-ray energy for 10 seconds.

### 3-point bending strength test

3 point bending strength test was performed as previously described (Mohan et al., 2000). Tibiae were fixed in 10% formalin for 3-5 days at 4°C and stored frozen in gauze moistened in PBS with 0.01% sodium azide, prior to thawing in PBS at 4°C. samples were tested by three-point bending with the Instron DynaMight testing system (Model 8840; Instron, Canton, MA, USA).

### Bone histomorphometry

7-week-old mice were injected with calcein (20 mg/kg) at 8 days and 2 days before histomorphometric measurements on week 8 as described (Xing et al., 2013). Calcein retaining trabeculae and tartrate-resistant acid phosphatase (TRAP)-labeled trabecular surfaces were measured in a blinded fashion with the OsteoMeasure (OsteoMetrics, Decatur, GA, USA) software.

### Histology

Mouse femurs were fixed in 10% formalin overnight, washed in PBS, decalcified in 10% EDTA (pH 7.4) at 4°C for 7 days while shaking, and embedded in paraffin for sectioning. Longitudinal sections of distal femurs were stained with alizarin red, and hematoxylin & eosin using standard procedures. Fracture calli were stained with Safranin O or acid phosphatase 5, tartrate resistant/TRAP (Sigma-Aldrich) followed by fast green counterstain. TRAP (S387A, Sigma-Aldrich), Alizarin red (A5533, Sigma-Aldrich) and Alkaline phosphatase, ALP (N6125 & F3381, Sigma-Aldrich) staining of cell cultures were performed by standard procedures.

### Immunofluorescence

Longitudinal paraffin-embedded sections were processed as described (Gomez et al., 2022) following 1-hour antigen retrieval with 2mg/ml Hyaluronidase (Sigma-Aldrich) at 37°C. Sections were blocked in 2.5% normal horse serum and incubated overnight with primary antibodies for COL10A1 at 1:100 (ab58632, Abcam), IBSP at 1:100 (gift from Dr. Renny Franceschi, University of Michigan), SPP1 at 1:300 (ENH094-FP, Kerafast), SP7/OSX at 1:100 (Ab22552, Abcam), KSR2 at 1:100 (nbp1-83553, Novus Biologicals). Protein expression was detected by species-specific secondary antibodies (VECTOR labs, DI-3088, and DI-1794), followed by DAPI (D1306, Invitrogen) counterstain before imaging.

### Microscopy

Epifluorescence images were obtained on a Leica Digital Microscope DMI6000B with Leica Applicate Suite X software or an Olympus FV3000 confocal microscope via FV31S-SW software. Colorimetric histological images were obtained with an Olympus DP72 camera attached to an Olympus DP72 camera through DP2-BSW software.

### Elisa

Serum levels of P1NP, and collagen type 1 C-terminal telopeptide (Ctx-1) EIA kits, all from Immunodiagnostic Systems (Gaithersburg, MD, USA) were obtained according to the manufacturer’s instructions. Serum alkaline phosphatase (ALP) activity was measured in samples diluted to 1:20 as reported (Xing et al., 2013)

### Western Blot

Immunoblots were processed by standard procedures. Cells were lysed in RIPA buffer with 1mM DTT, 1X protease inhibitor, and 1X phosphatase inhibitor cocktail (Sigma-Aldrich). Protein lysate concentrations were determined with a BCA protein assay (ThermoScientific) and 10μg of each lysate was boiled in 4X SDS dye, then loaded on 10% SDS-PAGE gels for immunoblotting on PVDF membranes. Membranes were blocked in 4% BSA in 1X TBS and probed with S6 ribosomal protein 1:1000 (2217, Cell Signaling Technologies), phosphor-S6 ribosomal protein 1:1000 (2215, Cell Signaling Technologies), or B-Actin 1:5000 (A1978, Sigma-Aldrich). Primary antibodies were detected with Goat anti-Rabbit IgG-HRP (A9169, Sigma) or Rabbit angi-Mouse IgG-HRP (NBB720-H, Novus Biologicals) at 1:15,000. Blots were detected with Immobilon Chemiluminescent HRP substrate (P90720, MilliporeSigma), and exposed on autoradiography film (1968-3057, USA Scientific). Scanned images were quantified on ImageJ software.

### Real time quantitative PCR

RNA was extracted from adipocyte depots, bones, or cultured cells with TRI reagent (Molecular Research Center INC, TR118) according to the manufacturer’s instructions, and purified on silica columns with E.Z.N.A. Total RNA Kit I (R6834-02, Omega BIO-TEK). Total RNA was reverse transcribed to cDNA with Oligo(dT)12-18 and Superscript IV Reverse transcriptase (18091050, Invitrogen). Real time PCR reactions were processed on a ViiA 7 RT-PCR system (Applied Biosystems). All reactions were standardized with peptidyl prolyl isomerase A (*Ppia*) primers. Primer sequences used for RT-qPCR are listed in Supplementary file 1. Fold changes were calculated by the Delta Ct method.

### Cell culture

All cells were maintained in standard normoxic conditions; humidified, 37°C, 5% CO_2_ with 1% Penicillin/Streptomycin (Gibco). ST2 stem/stromal cell line was obtained from the American Type Culture Collection (Manassas, VA), tested negative for mycoplasma, and were authenticated by their ability to differentiate into chondrocytic, adipocytic, and osteoblastic lineages in their respective differentiation media. For gain of function studies, the coding region of GFP in pRRLin-CPPT-SFFV-E2A-GFP-wpre (LV-GFP) was swapped with that of *Ksr2* from pcDNA3-*Ksr2*-flag (Addgene), producing pRRLsin-CPPT-SFFV-E2A-*KSR2*-wpre (LV-KSR2). Lentivirus (LV) plasmids were co-transfected with Pax2 and VSVG plasmids in 293T cells for LV generation as previously reported (Lindsey, Xing, et al., 2019). LV particles were transduced directly into ST2 or BMSCs. ST2 cells were cultured in 10% CS (Hyclone) with no ascorbic acid (Life Technologies). Osteoblast differentiation was performed with 10mM Beta-glycerophosphate, BGP, and 50μg/mL ascorbic acid, AA (Sigma-Aldrich), with BGP only serving as vehicle. Glucose, insulin, rapamycin, and IGF-1 (MilliporeSigma) were added at concentrations mentioned in the text, and low glucose (LG) was 5.5mM, while high glucose (HG) was 25mM. Mission lentiviral transduction shRNA particles for control (SHC002V) and *Ksr2* (TRCN0000378606) were obtained from MilliporeSigma, and cells were selected in 10μg/mL puromycin for 1 week before osteoblast differentiation.

Ex vivo culture of calvarial osteoblasts were isolated from 21-day old C57BL/6J mice and maintained in 10% FBS (Gibco) AMEM no ascorbic acid (Life Technologies), before osteoblast differentiation. Bone marrow stem/stromal cells (BMSCs) were isolated from whole femurs and tibias of 4- to 6-week-old C57BL/6J mice, while macrophages were isolated from 8-week-old C57BL/7J mice. KSR2 and GFP were overexpressed by lentivirus, without antibiotic selection. Osteoclast differentiation was performed with 30ng/mL MCSF (R&D), and 30ng/mL RANKL (R&D), with MCSF only serving as vehicle controls.

### Figures

Figures were assembled on Adobe Illustrator CS5. Quantitative graphs were generated on PRISM v9.3.1 software (Graphpad).

### Statistics

Statistical analysis was performed by two-tailed student t-test on Excel (Microsoft Office 365) following tests for normality. Data are presented as mean ± standard error of the mean (SEM) throughout. Values were considered significant at P < 0.05 or less

### Study Approval

Animal studies were performed according to protocols approved by the Institutional Animal Care and Use Committee of the VALLHS.

## Author contributions

SM Conceived the project. SM, GAG, CK, WX, CHR, RCL and DRP contributed to methodology. GAG, SP, CK, WX and CHR performed experiments. GAG, SP, and CHR performed data analysis. GAG, SM wrote and edited the original draft of the manuscript. RCL, DRP, WX, CHR, and CK reviewed and edited the manuscript. SM acquired the funding and supervised the study.

## Acknowledgements

We would like to thank Dr. Robert Brommage for helpful critical comments on the manuscript; Dr. Andrew S. Shaw and Dr. McMahon for mice, and Dr. Renny Franceschi for the BSP antibody; and Jasmine Lau, Fern Baedyananda, William Tambunan, Destiney Larkin, Nancy Lowen for excellent technical assistance. This work was supported by grants from National Institutes of Health (R01 AR048139 and R21 AG062866) and U.S. Department of Veterans Affairs (101 BX005263 and Ik6 BX005381) awarded to SM. SM is a recipient of Senior Research Career Scientist Award U.S. Department of Veterans Affairs.

---

**Figure 3—figure supplement 1.**
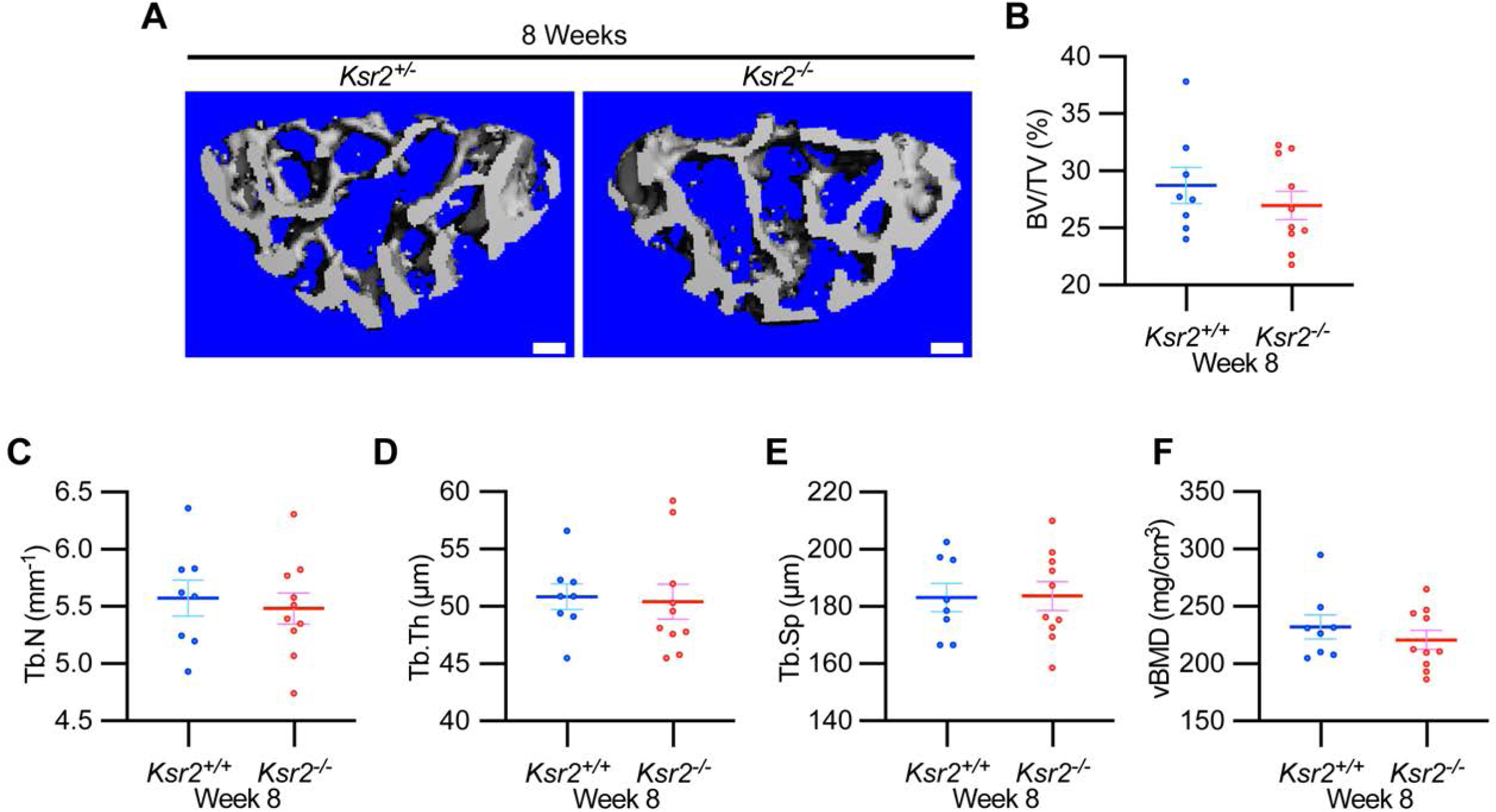
Vertebral Trabecular bone is not affected by Ksr2. (A) Representative 3D microCT reconstruction images of lumbar vertebrae in wild-type (*Ksr2^+/+^*, WT) or knockout (*Ksr2^-/-^*, KO) mice at 8 weeks, revealing no change in trabecular bones by *Ksr2* deletion. Scale bar: 100μm. (B-F) MicroCT measurements from the trabecular bone as represented in part A (n = 8-10/group), mixed genders. Abbreviations: BV/TV, Bone Volume/Tissue Volume; Tb.N, Trabecular number; Tb.Th, Trabecular thickness; Tb.Sp, Trabecular spacing; vBMD, volumetric Bone mineral density. Statistics analyzed by unpaired 2- tailed student t-test, and graphed lines represent the mean ± SEM.

**Figure 3—figure supplement 2.**
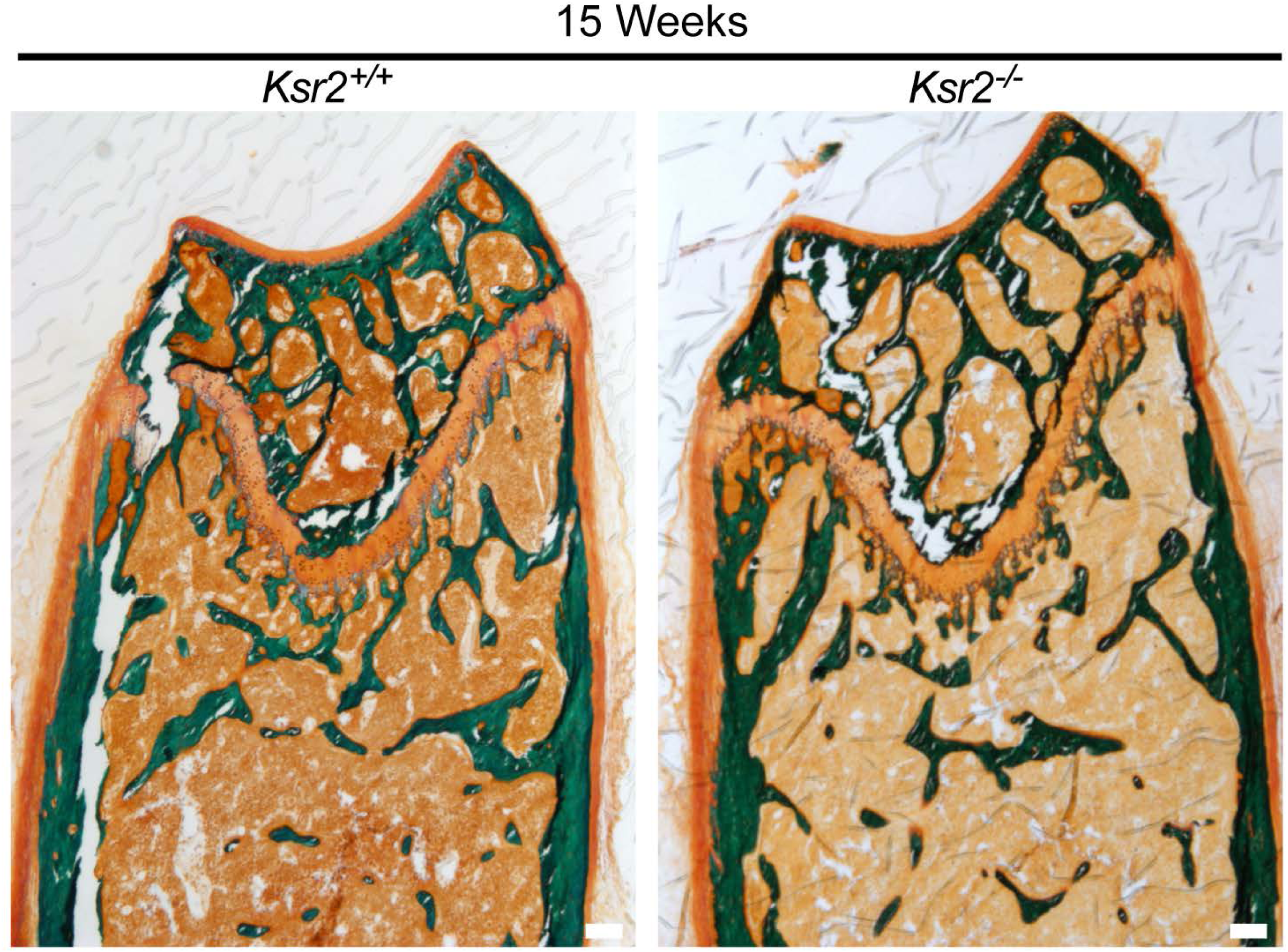
Osteoid area is regulated by Ksr2. Representative Goldner stained longitudinal sections of distal femoral bones used to quantify osteoid area over bone area (OA/BA) in Figure 3J. Scale bar: 100μm

***List of source data files***

**Titles.** Legends

***(Figure 1- Source data 1)***

***Source Data for Figure 1C-1H.*** microCT measurements of female trabecular bone.

***(Figure 1- Source data 2)***

**Source Data for Figure 1J-1M.** microCT measurements of female primary spongiosa.

***(Figure 1- Source data 3)***

**Source Data for Figure 1O-1Q.** microCT measurements of female cortical bone.

***(Figure 2- Source data 1)***

**Source Data for Figure 2B-2G.** microCT measurements of male trabecular bone.

***(Figure 2- Source data 2)***

**Source Data for Figure 2I-2K.** microCT measurements of male cortical bone.

***(Figure 2- Source data 3)***

**Source Data for Figure 2L-2Q.** microCT measurements of Ksr1 knockout mice.

***(Figure 3- Source data 1)***

**Source Data for Figure 3B-3G.** X-ray measurements of Ksr2 knockouts in DBA/1LacJ.

***(Figure 3- Source data 2)***

**Source Data for Figure 3I-3J.** Distal femur-Alizarin red quantification.

***(Figure 3- Source data 3)***

**Source Data for Figure 3L-3N.** Histomorphometric measurements of Ksr2 knockout mice.

***(Figure 3- Source data 4)***

**Source Data for Figure 3O-3Q.** Serum Elisa measurements of Ksr2 knockout mice.

***(Figure 3- Source data 5)***

**Source Data for Figure 3T.** RT-qPCR data of 12 week old Ksr2 knockout femurs versus wild type. Average fold changes plotted and T-test values are highlighted in yellow.

***(Figure 3- figure supplement 1- Source data 1)***

**Source Data for Figure 9- Supplement 1.** RT-qPCR data for RUNX2 shRNA. Average fold changes plotted and T-test values are highlighted in yellow.

***(Figure 4- Source data 1)***

**Source Data for Figure 4B-4C.** RT-qPCR data of 28 week old Ksr2 knockout versus wild type (adipose tissue). Average fold changes plotted and T-test values are highlighted in yellow.

***(Figure 4- Source data 2)***

**Source Data for Figure 4E.** Quantification of Osmium Tetroxide labeled microCT of tibia.

***(Figure 4- Source data 3)***

**Source Data for Figure 4G.** Quantification of Adipocytes from H&E stained femurs metaphysis.

***(Figure 4- Source data 4)***

**Source Data for Figure 4H.** RT-qPCR data of 12 week old Ksr2 knockout versus wild type femur (adipocyte markers). Average fold changes plotted and T-test values are highlighted in yellow.

***(Figure 4- Source data 5)***

**Source Data for Figure 4I.** RT-qPCR data of 12 week old Ksr2 knockout versus wild type femur (Wnt markers). Average fold changes plotted and T-test values are highlighted in yellow.

***(Figure 5- Source data 1)***

**Source Data for Figure 5D-5F.** microCT measurements of fracture callus after 3 weeks.

***(Figure 5- Source data 2)***

**Source Data for Figure 5H, 5J.** Quantification of Safranin O and ACP5/TRAP of fracture callus after 3 weeks.

***(Figure 5- Source data 3)***

**Source Data for Figure 5L-5N.** Quantification of immunofluorescence images of fracture callus after 3 weeks.

***(Figure 5- Source data 4)***

**Source Data for Figure 5O-5Q.** Quantification of three point bending tests.

***(Figure 6- Source data 1)***

**Source Data for Figure 6B.** RT-qPCR data of ex vivo osteoblast differentiation time-course. Average fold changes plotted and T-test values are highlighted in yellow.

***(Figure 6- Source data 2)***

**Source Data for Figure 6C.** RT-qPCR data of ex vivo osteoclast differentiation time-course. Average fold changes plotted and T-test values are highlighted in yellow.

***(Figure 6- Source data 3)***

**Source Data for Figure 6E.** RT-qPCR data for osteoblast differentiation from BMSCs. Average fold changes plotted and T-test values are highlighted in yellow.

***(Figure 6- Source data 4)***

**Source Data for Figure 6F.** RT-qPCR data for osteoblast differentiation from BMSCs. Average fold changes plotted and T-test values are highlighted in yellow.

***(Figure 6- Source data 5)***

**Source Data for Figure 6H.** Quantification of osteoclasts differentiated from primary macrophages.

***(Figure 6- Source data 6)***

**Source Data for Figure 6I.** RT-qPCR data for osteoclast differentiation from primary macrophages. Average fold changes plotted and T-test values are highlighted in yellow.

***(Figure 7- Source data 1)***

**Source Data for Figure 7A-7D.** X-ray measurements of Ksr2 knockout mice after pair-feeding experiments, and microCT of femur metaphysis in pair-fed mice.

***(Figure 7- Source data 2)***

**Source Data for Figure 7F-7G.** X-ray measurements of osteoblast-specific Ksr2-conditional knockout mice.

***(Figure 7- Source data 3)***

**Source Data for Figure 7J-70.** microCT measurements of distal femoral metaphysis from osteoblast-specific Ksr2-conditional knockout mice.

***(Figure 8- Source data 1)***

**Source Data for Figure 8A.** RT-qPCR data for ST2 stromal cells following osteoblast differentiation in low glucose or high glucose and either no insulin or with insulin.

Average fold changes plotted and T-test values are highlighted in yellow.

***(Figure 8- Source data 2)***

**Source Data for Figure 8C.** Quantification of western blot data.

***(Figure 8- Source data 3)***

**Source Data for Figure 8E.** RT-qPCR data for ST2 stromal cells with Ksr2 shRNA vs Control shRNA, following osteoblast differentiation in absence or presence of rapamycin. Average fold changes plotted and T-test values are highlighted in yellow.

***(Figure 8- Source data 4)***

**Source Data for Figure 8F-8G.** RT-qPCR data for ST2 stromal cells with Ksr2 OE vs GFP, following osteoblast differentiation in absence or presence of rapamycin. Average fold changes plotted and T-test values are highlighted in yellow.

***(Figure 8- Source data 5)***

**Source Data for Figure 8H.** RT-qPCR data of 12 week old Ksr2 knockout versus wild type femur (Hypoxia and Notch pathway related genes). Average fold changes plotted and T-test values are highlighted in yellow.

***(Supplementary File 1)***

**Methods Table1.** Real Time quantitative PCR Primers used in this study

***(Supplementary File 2)***

**Western blot.** Images of western blot film used in Figure 8B. Note that membranes probed for RPS6 and p-RPS6 were exposed simultaneously, as were corresponding membranes probed with B-ACTIN.

